# Structural basis of Ty1 integrase tethering to RNA polymerase III for targeted retrotransposon integration

**DOI:** 10.1101/2022.03.07.483246

**Authors:** Phong Quoc Nguyen, Sonia Huecas, Amna Asif-Laidin, Adrián Plaza-Pegueroles, Noé Palmic, Joël Acker, Juan Reguera, Pascale Lesage, Carlos Fernández-Tornero

## Abstract

The yeast Ty1 retrotransposon integrates upstream of genes transcribed by RNA polymerase III (Pol III). Specificity of integration is mediated by an interaction between the Ty1 integrase (IN1) and Pol III, currently uncharacterized at the atomic level. Here, we report cryo-EM structures of Pol III in complex with IN1, revealing a 16-residue segment at the IN1 C-terminus that contacts Pol III subunits AC40 and AC19, an interaction that we validate by *in vivo* mutational analysis. Unexpectedly, IN1 binding associates with insertion of subunit C11 C-terminal Zn-ribbon into the Pol III funnel, which provides atomic evidence for a two-metal mechanism during RNA cleavage. Moreover, unstructured regions of subunits C53 and C37 reorganize close to C11, likely explaining the connection between the C37/C53 heterodimer and C11 during transcription reinitiation. Our results suggest that IN1 binding induces a Pol III configuration that favors chromatin residence, thus improving the likelihood of Ty1 integration.

## Introduction

Most living organisms harbor transposable elements (TE) that, upon mobilization within the genome, participate in the adaptative response to environmental changes^1^. As mobile genetic elements, TEs also represent a potential threat for genome integrity and, accordingly, are at the ground of human pathologies including cancer and aging^2–4^. To replicate and yet minimize genetic damage to its host, TEs have evolved the capacity of integrating into specific regions of the genome with minimal effects on cell function, generally through tethering to different cellular machineries that operate on the DNA^5^. Long-terminal repeat (LTR) retrotransposons constitute a group of TEs that, like retroviruses, replicate by reverse transcription of their mRNA in a double-stranded DNA (cDNA) that is subsequently integrated into the genome by their own integrase. Ty1 is the most active and abundant LTR-retrotransposon in *Saccharomyces cerevisiae*^6,7^. Ty1 preferentially integrates into nucleosomes within the first kilobase upstream of genes transcribed by RNA polymerase III (Pol III)^8,9^.

Pol III transcribes short genes encoding for untranslated RNAs such as transfer RNAs (tRNA), the 5S ribosomal RNA (rRNA) and the U6 spliceosomal RNA. Pol III is constituted by 17 subunits that are organized into four architectural units^10–12^. The enzymatic core comprises Pol III-specific subunits C160 and C128 forming the DNA-binding cleft with the active site, the assembly AC40/AC19 heterodimer shared with Pol I, peripheral subunits Rpb5, Rpb6, Rpb8, Rpb10 and Rpb12 shared with RNA polymerase I (Pol I) and RNA polymerase II (Pol II), and subunit C11 involved in transcriptional pausing, RNA cleavage, termination and reinitiation^13–15^. The three remaining units are formed by Pol III-specific subunits: (i) a stalk including subunits C25 and C17, (ii) a nearby C82/C34/C31 heterotrimer sharing homology with TFIIE, and (iii) a TFIIF-like heterodimer formed by subunits C37 and C53, which cooperate with C11 in termination and reinitiation^14^.

Genome-safe Ty1 integration upstream of Pol III-transcribed genes is mediated by an interaction between the Ty1 integrase (IN1) and the Pol III AC40 subunit^16,17^. Contact with additional Pol III subunits including C53, C34 and C31 has also been reported *in vitro*^18^. The IN1 N-terminal half retains the well-structured, phylogenetically-conserved oligomerization and catalytic domains of retroviral integrases, while the less-conserved C-terminal half is intrinsically disordered^19^. We recently showed that a stretch of residues next to the C-terminal end of IN1 is necessary and sufficient for association with AC40, IN1 recruitment to Pol III-transcribed genes and Pol III-mediated integration^16^. Nonetheless, the molecular details of this interaction have been obscured by lack of structural studies.

## Results

### Cryo-EM structures of Pol III-IN1 complexes

To gain structural insights into IN1 tethering on Pol III, we first formed an *in vitro* complex between recombinant IN1 produced in bacteria^19^ and the Pol III enzyme isolated directly from yeast^20^. The two species interacted in a 1:1 stoichiometry as shown by native gel electrophoresis (Figure S1A), demonstrating a direct interaction between the two enzymes. This interaction can be stabilized through mild crosslinking, despite the appearance of higher order oligomers in minor extent. Using this sample, we reconstructed the cryo-EM structure of the Pol III-IN1 complex at 2.6 Å resolution (Figure 1A; Figure S1B-D, map A; Table S1). Besides, we formed a Pol III elongation complex (EC) using a transcription bubble mimetic comprising an 11-nucleotide mismatched DNA region, where one of the mismatched strands is hybridized to a 10-nucleotide RNA molecule (Figure 1, inset). Interestingly, IN1 is capable of interacting with this Pol III EC in an equivalent manner as with free Pol III (Figure S2A), indicating that the interaction occurs at the periphery of this multi-protein enzyme. This preparation was employed to obtain the cryo-EM structure of Pol III EC in complex with IN1 at 3.1 Å resolution (Figure 1B; Figure S2B-D, map D; Table S1). The structures of Pol III-IN1 in the absence and in the presence of nucleic acids show that Pol III adopts a global configuration that is equivalent to those of free Pol III in the closed state (Figure S3A) and the canonical Pol III EC (Figure S3B), respectively^12^. Nevertheless, despite overall similarities, the cryo-EM maps in the presence of IN1 exhibit remarkable differences to reported structures, as described below.

**Figure 1.**
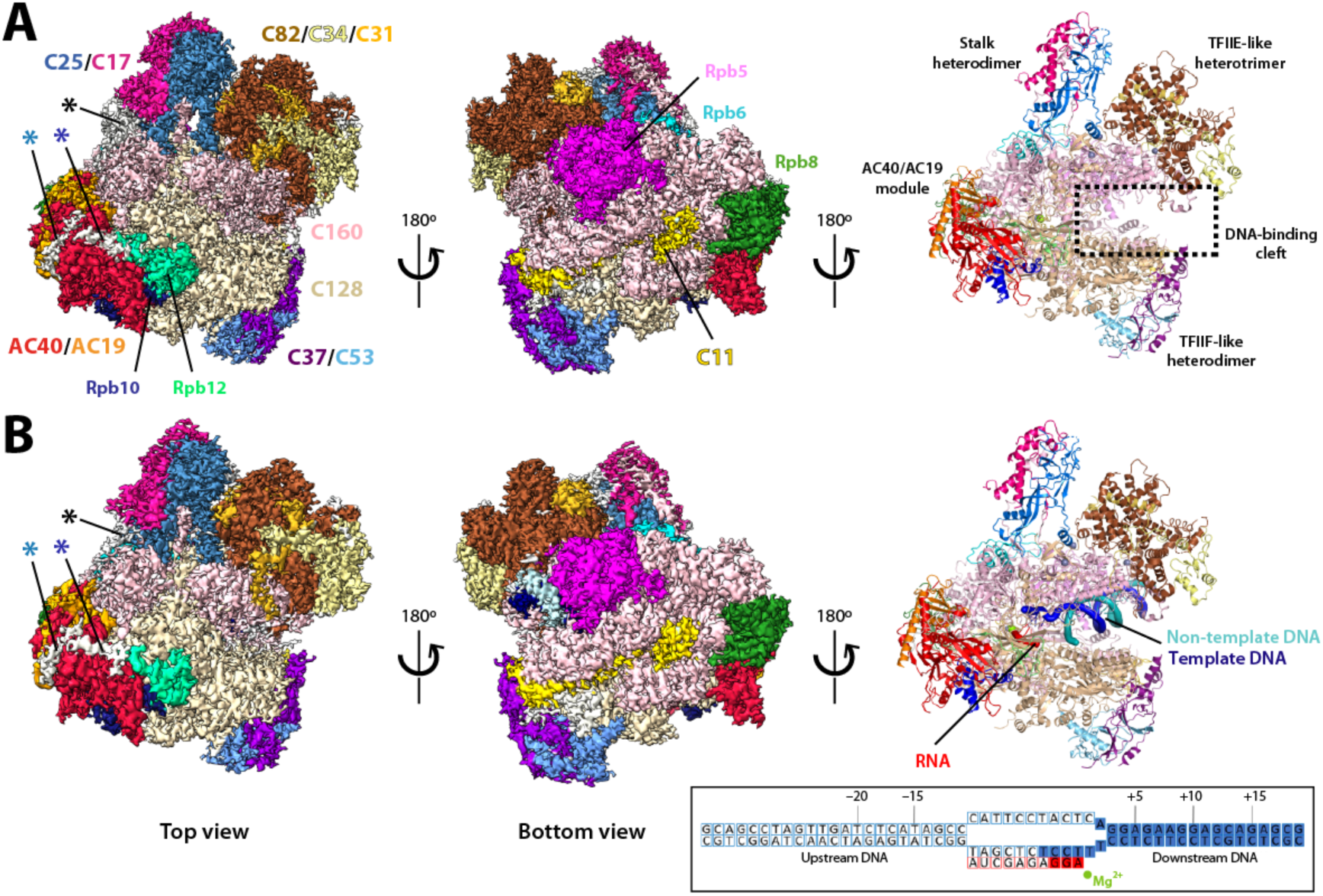
Overall structures of Pol III in complex with IN1. Cryo-EM map (left and middle) and resulting model (right) of Pol III-IN1 in the absence (A) or in the presence (B) of a mismatched transcription bubble (inset). All subunits and the main structural elements of Pol III are labelled. Blue, dark blue and black stars indicate three pieces of density in the vicinity of the AC40 subunit. Filled squares in the inset correspond to modelled nucleotides.

Both maps reveal two pieces of elongated density at the surface of subunit AC40, each located on one side of a hairpin-like loop (residues 108-130) of this subunit (Figure 1, light and dark blue stars). We unambiguously assigned one of these densities in the vicinity of AC40 to an IN1 segment (residues 609-625; Figure 2A, blue) proximal to the IN1 C-terminus (Figure 2B, top). This segment contains the reported tethering motif of IN1 involved in Ty1 integration targeting (residues 617-622), identified through mutational analysis and referred to as TD1^16^. Our structures show that this motif is extended to IN1 residues 609-625 and, thus, we will hereafter employ TD1 to designate the whole AC40-binding segment. TD1 is flanked by two nuclear localization signals (NLS; residues 596-598 and 628-630) defining a bipartite NLS^21,22^ (Figure 2B). In our maps both NLS are disordered, consistent with the idea that TD1 can be presented to Pol III by NLS-bound importin-*α* after nuclear import^16^. Moreover, conservation of TD1 in Ty2 and Ty4 retrotransposon integrases (Figure 2B), suggests that an equivalent mechanism may operate for their interaction with AC40^17^ and their preferential integration upstream of Pol III-transcribed genes^7^. The second piece of elongated density (Figure 2A, dark blue) was assigned to a secondary IN1 segment (residues 565-571) that is proximal to TD1 and exhibits conservation with Ty2 and Ty4 integrases (Figure 2B, bottom). This additional contact between AC40 and IN1 could reinforce the primary binding site.

**Figure 2.**
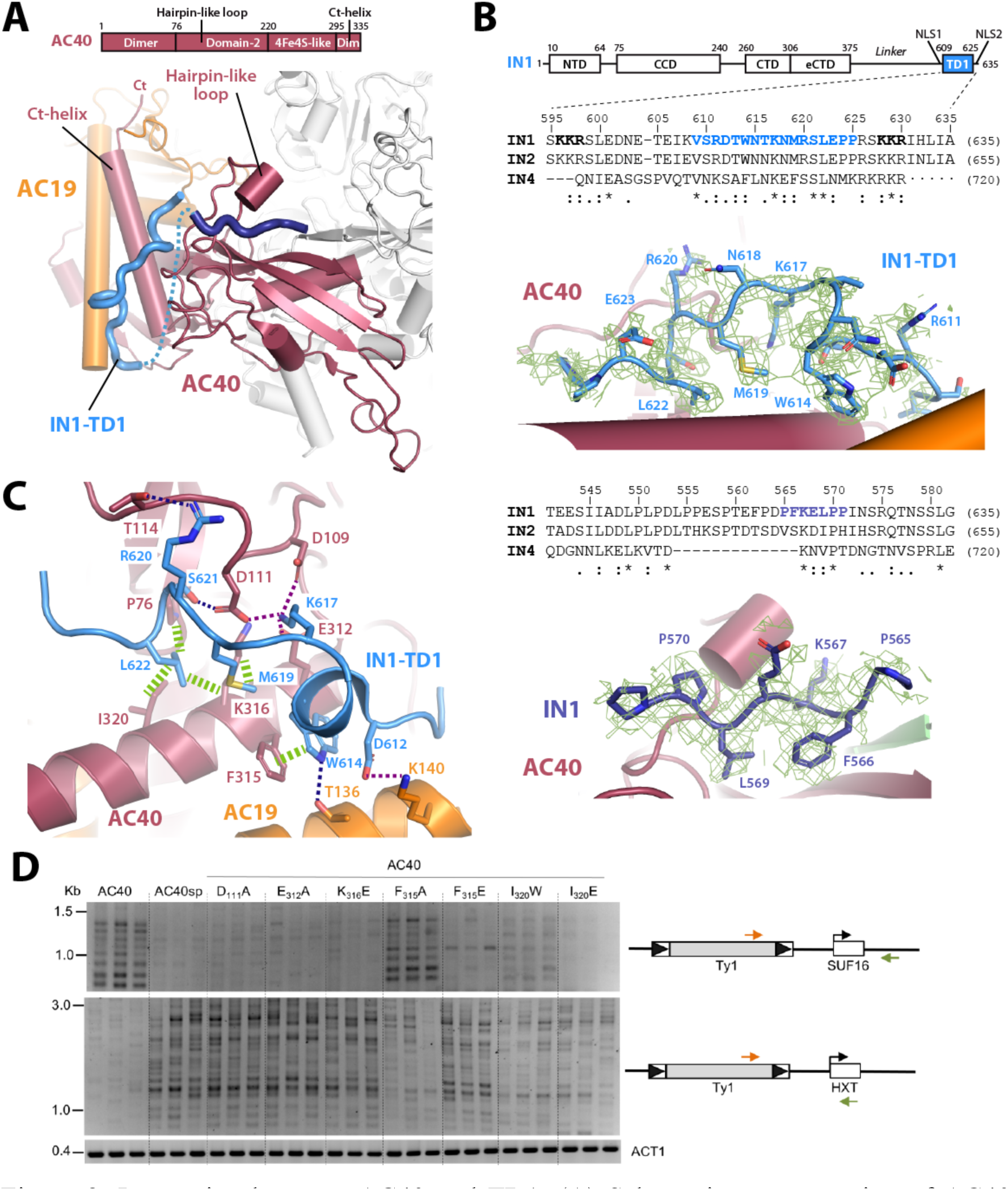
Interaction between AC40 and TD1. (A) Schematic representation of AC40 structural domains and close-up view of the Pol III-IN1 structure around AC40. Fully-modelled TD1 is shown in blue while a nearby peptide assigned to IN1 is in dark blue. (B) Schematic representation of IN1 structural domains including local alignments with Ty2 and Ty4 integrases (IN2 and IN4) around TD1 and the secondary binding site, with close-up view of the model (blue) and corresponding map (green mesh) around the two binding regions. Hyphens and interpuncts in the IN4 sequence correspond to gaps and insertions, respectively. (C) Atomic details of the AC40-IN1 interaction. H-bond, salt bridges and hydrophobic interactions are indicated with blue, purple and green dotted lines, respectively. (D) Detection of endogenous Ty1 insertions upstream of the *SUF16* Pol III-transcribed gene and the *HXT* subtelomeric genes (*HXT13, HXT15, HXT16* and *HXT17*) by PCR. Endogenous Ty1 retrotransposition was induced in cells growing at 20°C during 3 days in YPD media. Total genomic DNA was extracted from three independent cultures. *ACT1* is genomic DNA quality control.

A third piece of additional density (Figure 1, black star) exhibits a globular shape and localizes next to the Pol III stalk, both contacting a peripheral loop (residues 35-51) of subunit C17 and an *α*-helix (residues 15-20) in the tip domain of subunit C25 (Figure 3A). This density also contacts the nearby dock domain of subunit C160. Focused 3D classification around this density shows significant conformational heterogeneity (Figure 3B), thus hampering atomic modelling of this map region. Nevertheless, the absence of proximal flexible Pol III regions suggests that this density belongs to IN1. Superposition of our structures with that of Pol III in the presence of TFIIIB^23–25^ shows that IN1 binding to Pol III is compatible with the formation of the pre-initiation complex (Figure 3C). However, the third piece of globular density lies next to the N-terminal Zn-ribbon in the Brf1 subunit of TFIIIB, suggesting an interaction with IN1 that may retain TFIIIB on promoter DNA. As TFIIIB complexes cover up to 60 base pairs upstream of Pol III genes^26^, this is coherent with the absence of Ty1 integration within the first 80 base pairs upstream of tRNA genes^8,9^. Besides, superposition of our structures with that of the Pol I pre-initiation complex (PIC)^27^ shows that the position of the third piece of globular density is occupied by Rrn3 (Figure 3D), involved in Pol I activation prior to promoter recruitment^28,29^. This is consistent with lack of Ty1 integration next to genes transcribed by Pol I, in spite of IN1 recruitment at these loci mediated by AC40 binding^16^.

**Figure 3.**
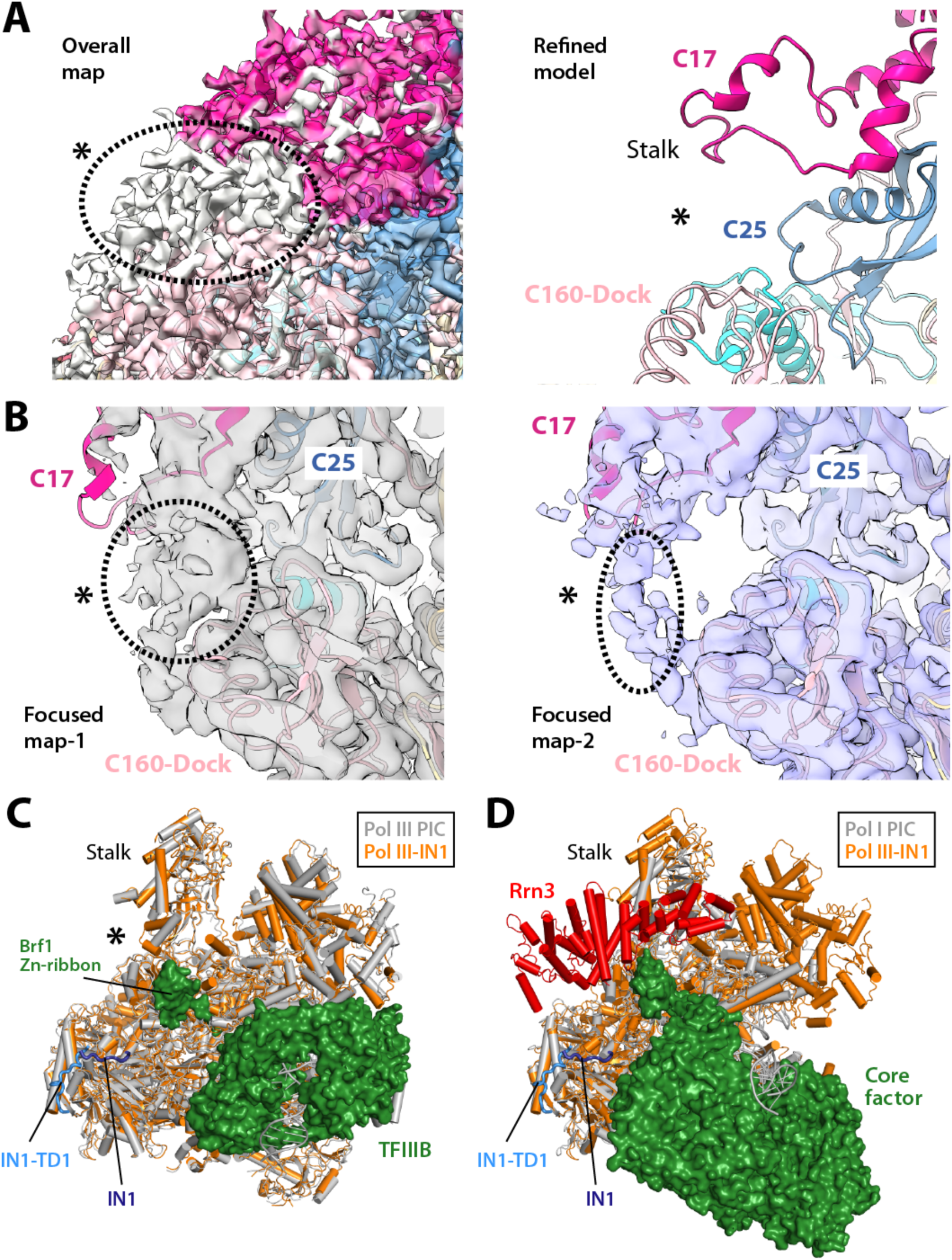
Structure of Pol III-IN1 around the Pol III stalk. (A) Pol III-IN1 map (left panel) and corresponding atomic model (right panel) around the Pol III stalk with a black star indicating additional density. (B) Two of the Pol III-IN1 maps and corresponding atomic models derived from subclassification using a mask around the Pol III stalk. (C) Superposition of the Pol III-IN1 structure (orange) and that of the Pol III pre-initiation complex (PIC) (PDB 6CND, gray for Pol III and green for TFIIIB). A black star indicates location of additional density in the Pol III-IN1 structure, as shown in panels A and B. (D) Superposition of the Pol III-IN1 structure (orange) and that of the Pol I PIC (PDB 6TPS, gray for Pol I and green for Core Factor). Activation factor Rrn3 is in red.

### Atomic details of the interaction between TD1 and AC40

Detailed analysis of the Pol III-IN1 complex revealed that TD1 binds a crevice on the surface of AC40 (Figure 2A) that is formed by the AC40 C-terminal *α*-helix, involved in heterodimerization with subunit AC19, and an AC40 hairpin-like loop (residues 108-130). TD1 adopts an extended conformation that includes a helical turn along the AC40 crevice and forms an intricate network of interactions with this subunit (Figure 2B-C). Residues W614, M619 and L622 in IN1 establish hydrophobic contacts with P76, F315, I320 and the aliphatic chain of K316 in AC40. In particular, the stacking interaction between W614 in IN1 and F315 in AC40 is expected to secure the interaction. Additionally, R620 and S621 in IN1 establish one hydrogen bond (H-bond) each with D111 and T114 in AC40, respectively. Importantly, K617 in IN1 appears as a central residue coordinating three salt bridges with D109, D111 and E312 in AC40. The Pol III-TD1 interaction is reinforced by contacts with subunit AC19, with residues T136 and K140 in this subunit respectively establish an H-bond and a salt bridge with W614 and D612 in IN1. Our structural observations correlate remarkably well with reported mutational analysis of IN1, showing that individual mutations in residues K617, S621 or L622 disrupt the AC40-IN1 interaction and alter Ty1 integration upstream of Pol III-transcribed genes^16^. This study also showed that mutation of residues N618 or E623, which in our structures point towards the solvent (Figure 2B), have no effect on the Pol III-IN1 interaction.

To investigate the role of individual AC40 residues at the tethering interface in Ty1 integration targeting, we produced single mutants at positions that appear critical according to our structures. Since AC40 is essential, we first checked that these mutations did not alter cell growth, AC40 protein levels or the integration frequency of a Ty1-his3AI reporter element expressed on plasmid from the *GAL1* promoter^30^ (Figure S4). For all mutants, we observed a less than twofold decrease in retrotransposition frequency, similar to that observed for the AC40 *Schizosaccharomyces pombe* ortholog (AC40sp), which behaves as a loss-of-interaction mutant^17^. Next, we performed qualitative PCR assay to monitor *in vivo* Ty1 insertion events at the *SUF16* tRNA gene and the HXT subtelomeric genes (Figure 2D). *SUF16* was identified as a hotspot of Ty1 integration in the presence of wild-type AC40, while the *HXT* loci were preferred target sites when the interaction was compromised using AC40sp or Ty1 elements mutated in IN1 targeting motif^16,17^. In comparison to the wild-type protein, AC40 mutants D111A, E312A or K316E induced a significant reduction of Ty1 integration events upstream of *SUF16* associated with a sharp increase in Ty1 integration at *HXT* loci, similar to the AC40sp mutant. An equivalent result was obtained when residue I320 in AC40 was mutated to either tryptophan or glutamate, suggesting that bulky or charged residues at this position are sufficient to disrupt IN1 tethering on Pol III. Notably, the I320W was designed because AC40sp contains a tryptophan at this position. Although to a lesser extent than I320E, I320W recapitulates the behavior of the AC40sp loss-of-function mutant. Additionally, mutation of F315 into alanine has no impact on Ty1 integration, while the F315E mutant behaves as the AC40sp mutant, highlighting the importance of the stacking interaction between this residue and W614 in TD1. In accordance, the W614A mutant in IN1, lacking in our former IN1 mutational analysis^16^, impaired the two-hybrid interaction of TD1 with AC40 (Figure S5A-B). This mutation also exhibited a reduction in integration events upstream of *SUF16* and an increase in Ty1 integration at *HXT* loci as compared to wild-type IN1, while it did not alter the overall Ty1 integration frequency (Figure S5C-D), indicating that this residue plays a major role in the Pol III-IN1 interaction. While alanine substitution of R620 in IN1 impaired the interaction with AC40^16^, this mutation displayed an intermediate phenotype with increased integration at subtelomeres despite a wild-type integration profile at *SUF16* (Figure S5C), suggesting a more complex role of R620 in Ty1 integration targeting.

Finally, we tried to rescue the AC40sp loss-of-function phenotype using single-residue mutagenesis. Superposition of the AC40sp structure^31^ onto our Pol III-IN1 structure shows that all residues of AC40 involved in TD1 binding are conserved with the exception of *S. cerevisiae* E312, which in *S. pombe* corresponds to V322 (Figure S6A-B). We produced the V322E mutation in the AC40sp loss-of-function mutant and performed *in vivo* Ty1 integration assays. We observed that, while the Ty1 integration pattern classically observed with wild-type AC40 at *SUF16* was not recovered, subtelomeric integration events were altered with relatively less integration compared to the AC40sp loss-of-function mutant (Figure S6C). This intermediate phenotype may reflect an improved interaction of AC40sp with IN1 while the contribution of the secondary binding sites is likely required to recover the wild-type phenotype.

### The C11 C-terminal Zn-ribbon inserts into the funnel pore

Subunit C11, involved in RNA cleavage^13^ as well as in termination and reinitiation together with the C37/C53 heterodimer^14^, comprises two Zn-ribbons (residues 1-36 and 69-110) connected by a flexible linker. In reported Pol III structures^12,23^, the C-terminal Zn-ribbon (C11-Ct) is either disordered or locates next to subunit Rpb5 (Figure S7A). Unexpectedly, our structures exhibit density in the Pol III funnel pore corresponding to C11-Ct (Figures 1 and 4A). While in recent structures of human Pol III EC this domain locates inside the funnel^32–34^, the human C11-Ct is retracted by about 9 Å from the active site respect to our structures (Figure S7B). This unanticipated location of C11-Ct suggests that IN1 binding might favor positioning of this domain in an RNA cleavage-prone configuration.

**Figure 4.**
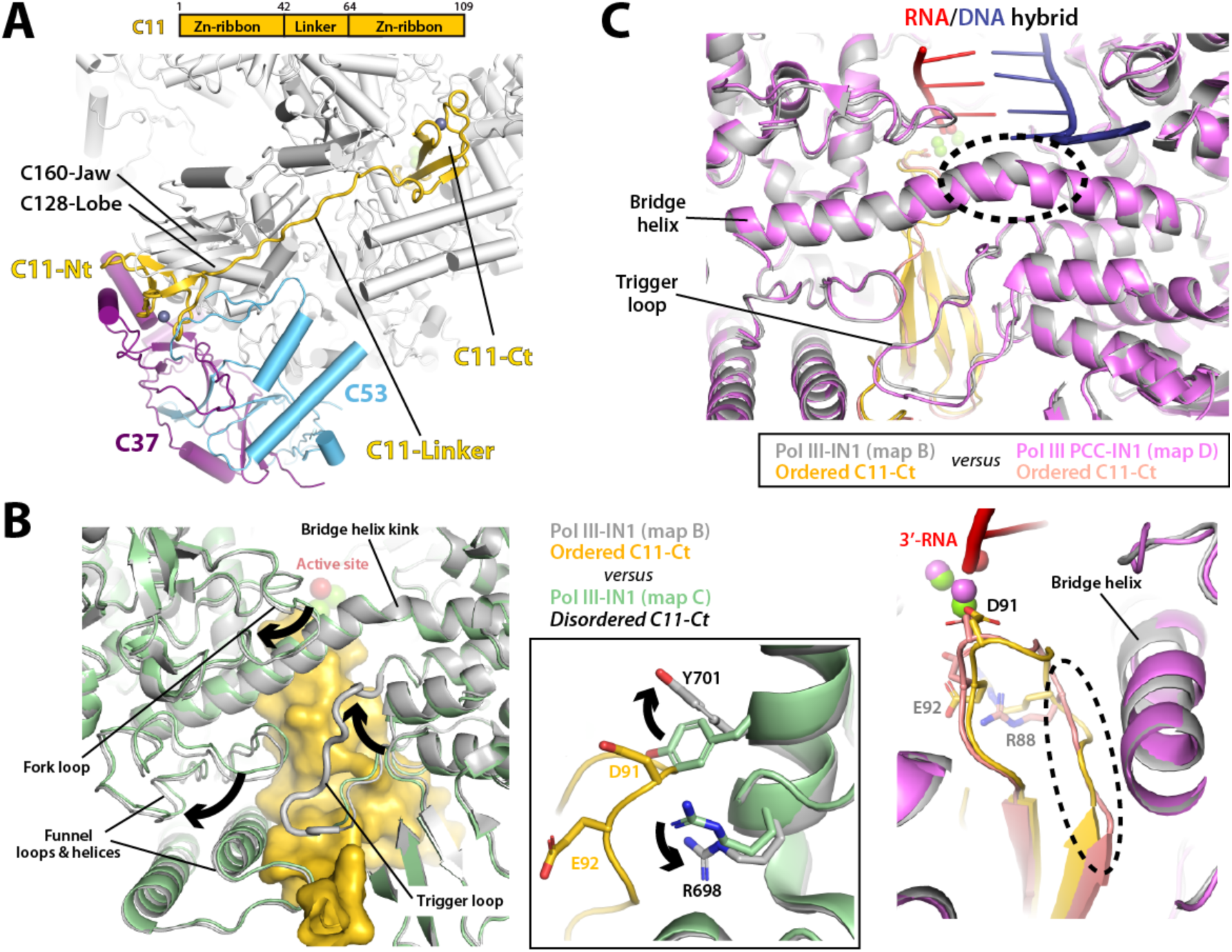
Structure of C11 in the presence of IN1. (A) Schematic representation of C11 and view around C11 of the Pol III EC-IN1 structure, representing a post-cleavage complex (PCC) of RNA incision. Subunits C37 and C53 are shown in purple and blue, respectively. (B) Structural comparison of the Pol III-IN1 models derived from subclassification using a mask around C11-Ct, showing presence (grey) and absence (green) of C11-Ct (yellow) in the Pol III funnel. Arrows indicate differences between the structures. The inset shows a close-up view around the C11 acidic loop and the gating tyrosine Y701 in subunit C128. (C) Structural superposition of Pol III-IN1 (grey) and Pol III PCC-IN1 (magenta) structures, with close-up views around the bridge helix (upper panel) and C11-Ct (lower panel). Dotted circles mark a kink in the bridge helix and remodeling of the C11 acidic loop, respectively.

Subclassification of the Pol III-IN1 dataset showed that approximately one third of the particles contain C11-Ct in the funnel (Figure S1, map B), while the remaining particles presented no density for this domain (Figure S1, map C). Comparison of these structures uncovered Pol III rearrangements for C11-Ct accommodation, mainly involving opening of the C160 funnel and C128 fork domains using a kink in the bridge helix as hinge, associated with ordering of the C160 trigger loop (Figure 4B). Additionally, residues R698 and Y701 in C128 switch conformation to avoid clashes with the C11 acidic loop (residues 88-93; Figure 4B, inset), involved in RNA cleavage^35^. Notably, Y701 corresponds to Y769 in the Rpb2 subunit of RNA polymerase II (Pol II), a residue proposed to limit the extent of backtracking^36^. Strikingly, subclassification of the dataset obtained in the presence of nucleic acids showed that virtually all particles contain C11-Ct in the funnel (Figure S2, map D). In this structure, the Pol III bridge helix presents a straight conformation, associated to a conformational change in the nearby C11 acidic loop, while the trigger loop remains ordered and retracted (Figure 4C). An equivalent configuration with a straight bridge helix and a locked trigger loop has been observed in the reactivated intermediate of backtracking for Pol II in complex with TFIIS^36^, the factor involved in RNA cleavage in this transcription system (Figure S7C, left panel).

The quality of our maps containing C11-Ct in the Pol III funnel (maps B and D) enabled precise building of all residues and metal ions in the active site (Figure 4A), where three catalytic aspartates (D511, D513, D515) from subunit C160 coordinate a Mg^2+^ ion (MgA in Figure 5B). In addition, residue D91 in C11-Ct together with catalytic residues D511 and D513 coordinate a second Mg^2+^ ion lying 3 Å away from MgA (MgB in Figure 5B), thus creating a composite active site with two metals as postulated for Pol II^36,37^. In the presence of nucleic acids, the RNA 3’ end further coordinates both Mg^2+^ ions (Figure 5B, right), while this coordination is replaced by a solvent molecule when nucleic acids are absent (Figure 5B, left). Superposition of the bacterial RNA polymerase structure in the pre-catalytic state^38^ shows that this water molecule lies 2 Å away from the phosphate next to the scissile bond and completes the tetrahedral coordination sphere of MgA (Figure 5C), suggesting that it might represent the nucleophilic water for RNA cleavage. Our structure in the presence of nucleic acids likely reflects a post-catalytic state of RNA cleavage that we term Pol III post-cleavage complex (Pol III PCC), equivalent to that observed for bacterial RNA polymerase^38^. Moreover, while the complete downstream dsDNA is observed in our map, density for the DNA/RNA hybrid only allows modelling of three base pairs next to the Pol III active site (Figure 5D). This is in sharp contrast with reported structures of canonical Pol III EC^12^, where eight base pairs are observed for the hybrid (Figure 5D), indicating that, in the presence of IN1, the interaction between Pol III and the DNA/RNA hybrid is destabilized. Altogether, these observations suggest that, in the presence of IN1, Pol III adopts a distinctive conformational state (Pol III PCC-IN1) that might influence Pol III function *in vivo*. In agreement, the *in vitro* Pol III transcriptional activity is mildly reduced in the presence of IN1 (Figure 5E). Our RNA extension assay shows that synthesis of intermediate RNA products is decreased over time when Pol III is preincubated with IN1, while the runoff product is less affected, suggesting a shorter dwell time at pausing sites for Pol III in the presence of IN1.

**Figure 5.**
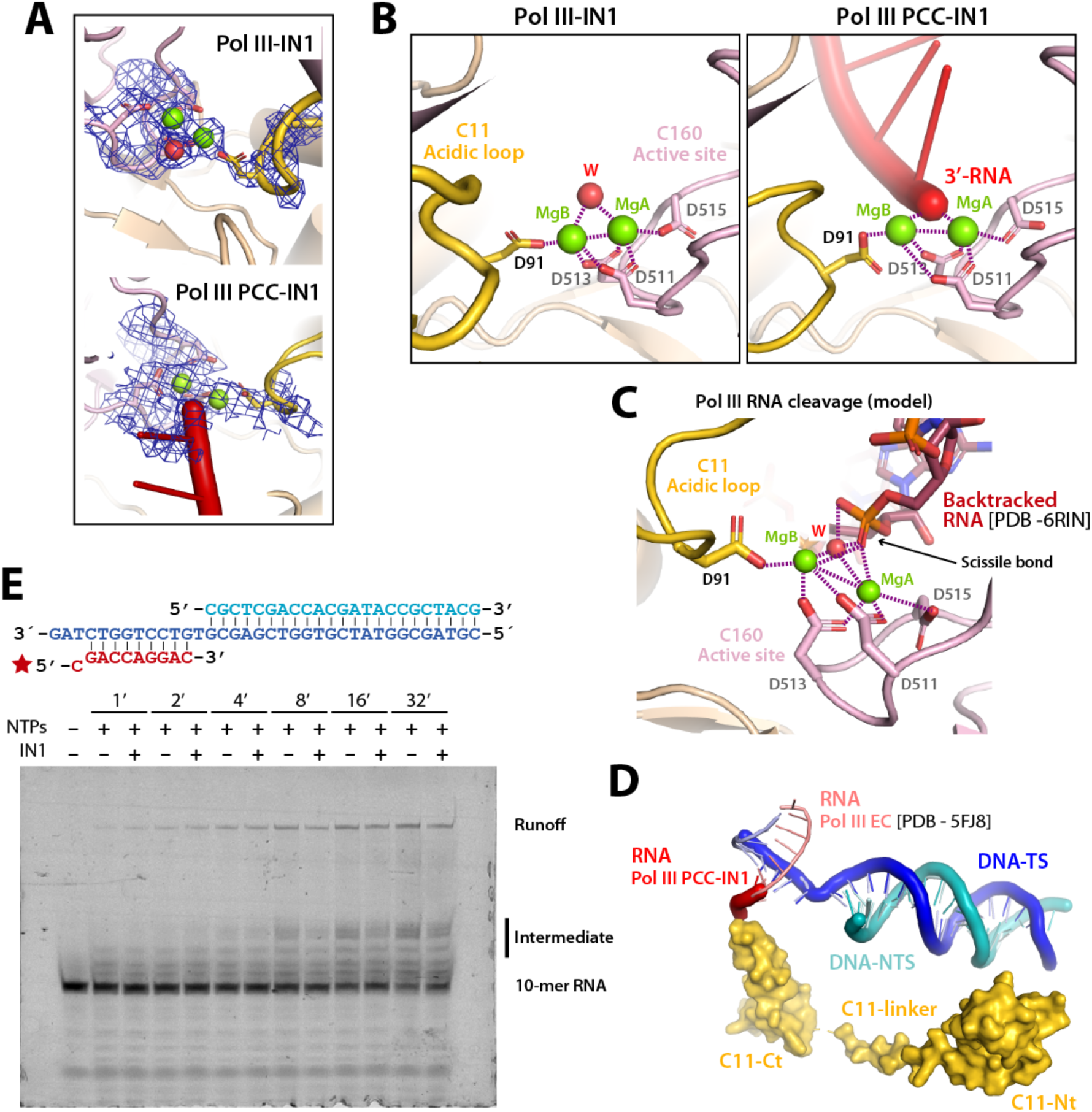
Pol III active site for RNA cleavage. (A) Active site model and map (blue mesh) in the structures of Pol III-IN1 (upper panel) and Pol III PCC-IN1 (lower panel). (B) Close-up views of the active site in the structures of Pol III-IN1 (left panel) and Pol III PCC-IN1 (right panel). Green and red spheres correspond to Mg^2+^ ions and water, respectively, while dotted lines indicate metal coordination. (C) Model of a pre-catalytic complex for RNA cleavage using our Pol III-IN1 structure and an RNA molecule from superposition with bacterial RNA polymerase in pre-catalytic state (PDB 6RIN). (D) Close-up view around the cleft of the superposition between the Pol III PCC-IN1 structure (thick ribbon nucleic acids) and that of the canonical Pol III-EC (PDB 5FJ8; thin ribbon nucleic acids). The C11 subunit in Pol III PCC-IN1 is shown as yellow surface, while the DNA template strand (TS) and non-template strand (NTS) are blue and cyan, respectively, and the RNA is red. (E) RNA extension assay of Pol III (100 nM) in the absence (-) or in the presence of IN1 (200 nM) at different time points. The nucleic acid scaffold employed in the assay is shown above the gel, with RNA in red and DNA in cyan and blue. A red star indicates a fluorescent label on the RNA 5’ end.

### The C11 N-terminal Zn-ribbon and linker associate with C37/C53

In our structures, the C11 N-terminal Zn-ribbon (C11-Nt) locates between the C160 jaw, the C128 lobe and subunit C37 (Figure 6A), in an equivalent position to that observed in Pol III structures in the absence of IN1^12^. We were able to model the entire C11 linker (residues 37-68) when C11-Ct is ordered (Figure S1, map B; Figure S2, map D), independent of the presence of nucleic acids. However, only the N-terminal third of the C11 linker (residues 37-47) is visible when the C11-Ct is disordered (Figure S1, map C). In our structures, this segment forms a β-strand extending a four-stranded β-sheet in the C160 jaw domain (Figure 6A). In contrast to published structures, ordering of this C11 linker region associates with the nearby appearance of elongated densities that further extend the jaw β-sheet (Figure 6A). These densities correspond to the N-terminal low-complexity region of C53^11^, where an *α*-helix followed by a long β-strand (residues 200-225) and another short β-strand become ordered next to the C11 linker. In agreement, the C11-Nt and linker are essential for Pol III reinitiation^15^ and C11 acts concertedly with the C37/C53 heterodimer during termination and reinitiation^14^, suggesting that Pol III might be more prone to reinitiate in the presence of IN1. Moreover, a potential role in Ty1 integration has been suggested for the low-complexity region of subunit C53^18^.

**Figure 6.**
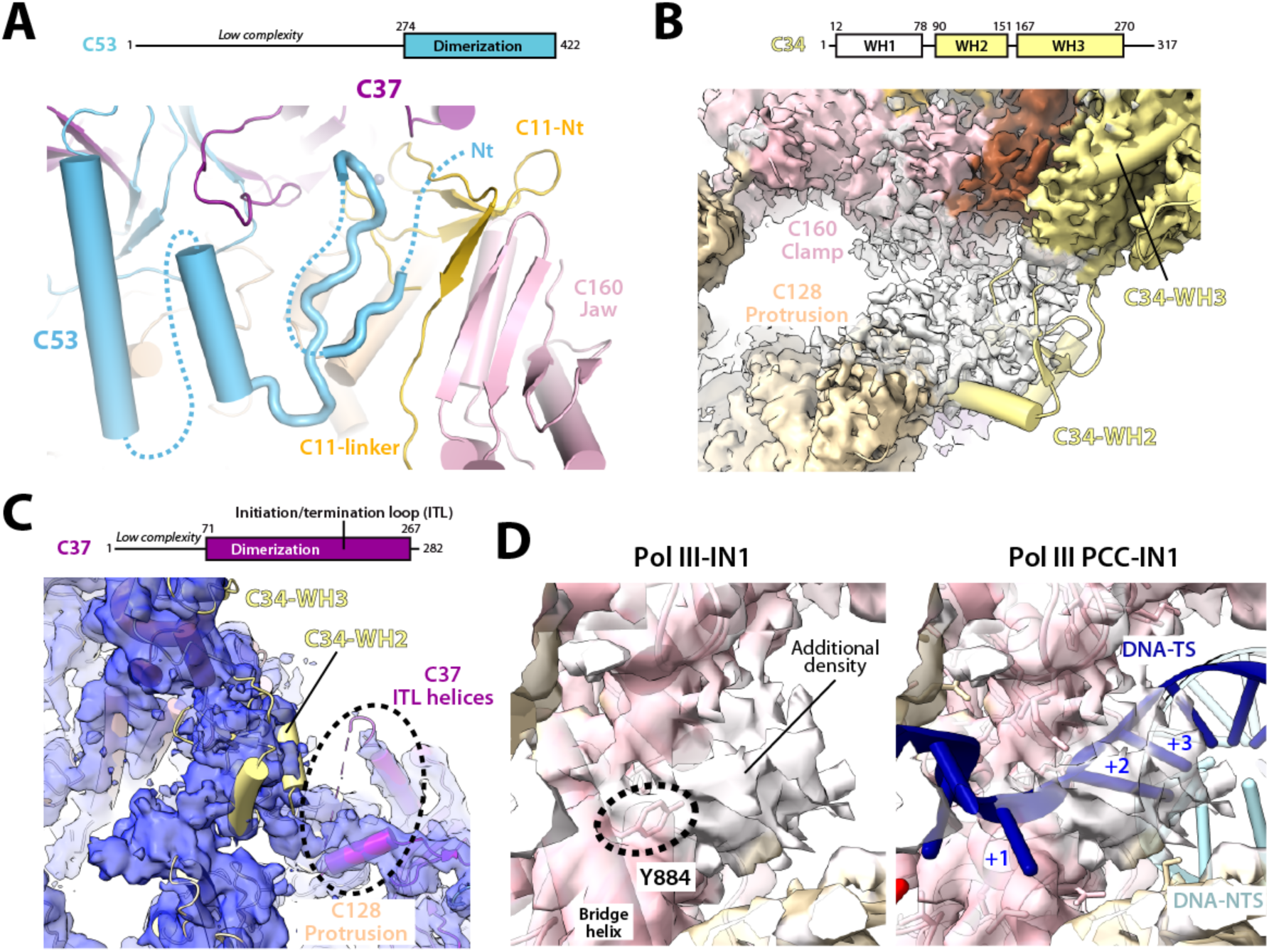
Structure of Pol III-specific structural elements in the presence of IN1. (A) Schematic representation of C53 and view of the Pol III-IN1 map around the C53 N-terminal region. (B) Schematic representation of C34 and view of the Pol III-IN1 map around C34, where the structure of Pol III from the reported Pol III-Maf1 complex (PDB 6TUT) has been fitted. (C) Schematic representation of C37 and view of the Pol III-IN1 map obtained from subclassification using a mask around C34, where the structure of Pol III from the reported Pol III-Maf1 complex (PDB 6TUT) has been fitted, showing partial ordering of the C37 initiation/termination loop (ITL) helices next to C34. (D) Close-up view of the Pol III-IN1 (left panel) Pol III PCC-IN1 (right panel) maps, with fitted model around the Pol III bridge helix. Nucleotides +1 (templating), +2 and +3 in the template strand (TS) are indicated.

### Partial ordering of the C34 WH2 domain at the rim of the cleft

The Pol III-IN1 map exhibits a piece of globular density at the rim of the DNA-binding cleft, between the C128 protrusion and the tip of the C160 clamp coiled-coil (Figure 6B). The dimensions of this density and its position next to the WH3 domain of subunit C34 led us to hypothesize that it corresponds to the WH2 domain (residues 89-157) of this subunit. In agreement, this domain occupies an equivalent position in the Pol III structure complexed to transcriptional repressor Maf1^39^ (Figure 6B). Focused 3D classification confirmed the presence of this density in about 15% of the particles, also showing an additional density next to C34-WH2 that might account for the WH1 domain of this subunit (Figure 6C). In our structures, the WH2 domain of C34 connects to the initiation/termination loop (residues 211–224) and neighboring helices of subunit C37 (Figure 6C), as observed in Pol III initiation intermediates where WH1 and WH2 domains of C34 are ordered^23–25^. These results suggest that, in the presence of IN1, the Pol III enzyme adopts a configuration that may favor interaction with TFIIIB, thus facilitating promoter recruitment or retention.

Finally, the Pol III-IN1 map in the absence of nucleic acids including all particles (map A) presents a globular piece of density next to the Pol III bridge helix residue Y884 in subunit C160 (Figure 6D, left). This density, which is absent in reported structures of free Pol III^12,23^, is occupied by nucleotides +2 and +3 from the template strand in the Pol III PCC-IN1 (Figure 6D, right). While assignment of this density is not obvious due to poor map quality in this region, it might be occupied by the N-terminus of the nearby C37 subunit, which is predicted disordered^11^. The presence of this density within the Pol III cleft is incompatible with nucleic acid binding within the cleft. An equivalent role in preventing nucleic acid binding has been assigned to the DNA-mimicking loop of Pol I^40^, which occupies the cleft in the hibernating state of this enzyme^28,41^.

## Discussion

Targeted DNA integration is crucial to maintain genome integrity. The Ty1 retrotransposon takes advantage of the Pol III enzyme to integrate upstream of Pol III-transcribed genes, located in genome regions devoid of essential genes, thus protecting the yeast genome. The structures reported here provide evidence of primary IN1 binding through its TD1 on Pol III subunits AC40 and AC19 (Figures 1 and 2). Our structural and mutational analyses allowed definition of the precise TD1 boundaries, comprising residues 609 to 625, and support that the TD1-AC40 interaction is sufficient for IN1 tethering on Pol III, while a secondary contact with AC40 reinforces the interaction (Figure 2A-B). Additionally, a putative IN1 motif next to the Pol III stalk is blocked by Rrn3 in Pol I transcription system while the primary and secondary sites are available (Figure 3), suggesting a regulatory role in Ty1 integration for this motif.

Our maps show no density for the structured N-terminal half of IN1, containing functional domains for integration of the Ty1 cDNA. This suggests that the ∼230 residue linker connecting the N-terminal half to the C-terminus of IN1 (Figure 2B), which is disordered^19^, provides significant flexibility between the Pol III-tethering and DNA-integration modules of IN1. Ty1 integration occurs within the first to the third nucleosomes upstream of Pol III and, thus, a flexible linker is advantageous to reach varying distances while keeping strong tethering on genome-attached Pol III. This is in sharp contrast with integrases from other LTR-retrotransposons or retroviruses, all lacking a large disordered linker connected to their respective targeting domain^5^. Retroviruses use transcriptional regulators of Pol II^42^, while the Ty3 integrase interacts with DNA-bound TFIIIB in a configuration that blocks Pol III binding, thus hampering transcription of the corresponding Pol III gene^43^.

We show that IN1 binding associates with reordering of different Pol III regions that are distant from the IN1 binding sites. Unexpectedly, C11-Ct locates within the Pol III funnel, with its acidic loop complementing and remodeling the Pol III active site (Figure 4). Notably, our structures show how a two-metal catalytic mechanism could operate for RNA cleavage, thus providing a model for this essential RNA polymerase activity (Figure 5B-C). While this configuration may interfere with the transcription process, it may as well reduce Pol III pausing and its eventual detachment from DNA (Figure 5E). In addition, the otherwise disordered N-terminal region of subunit C53 partially orders next to the C11 linker (Figure 6A), which likely provides structural evidence for Pol III reinitiation and recycling. Altogether, we speculate that IN1 binding induces a unique conformational state of Pol III that increases its overall residence on the chromatin without significantly affecting the overall RNA production by this enzyme^44^. This is expected to enhance chances for IN1 to establish productive integration complexes at upstream nucleosomes.

Besides, the two NLSs of IN1 flanking TD1 and allowing IN1 nuclear import by importin-α^21,22^ are disordered in our structures, which allows simultaneous binding of importin-α and Pol III by IN1^16^. Moreover, structural superposition with available structures shows no clash between IN1 and TFIIIB. These observations are consistent with the interaction between IN1 and Pol III taking place either in the nucleoplasm or while Pol III is bound on the DNA. Therefore, the TD1 motif not only provides strong binding but also an increased likelihood for the tethering interaction to occur. This is especially relevant for the design of new gene therapy vectors able to target safe regions of the genome. The structures reported here represent a valuable tool in this respect, as they allow atomic-level understanding of the molecular mechanisms underlying targeted DNA integration upstream of genes transcribed by Pol III.

## Methods

### Growth media, yeast strains, and plasmids construction

*Saccharomyces cerevisiae* strains used in this study were grown using standard methods and are listed in Table S2. Plasmids were constructed using standard molecular biology procedures. Mutations were introduced in plasmids with Q5^®^ Site-directed mutagenesis (NEB). All the constructs were validated by DNA sequencing (Eurofins Genomics). All plasmids and primers used in this study are reported in Tables S3 and S4, respectively.

### *RPC40* mutant strains construction

To construct *RPC40* mutants, we used a *S. cerevisiae* yeast strain (LV1690) deleted for its *RPC40* genomic copy (AC40sc) and transformed with a centromeric plasmid bearing *Schizosaccharomyces pombe RPC40* gene (pTET-HA-AC40sp, *URA3*) to ensure cell viability^17^. Mutations were introduced in *RPC40* carried on a centromeric plasmid (pTET-HA-AC40sc, *TRP1*). Yeast strain was transformed with mutant pTET-HA-AC40sc plasmids and cells were plated on DO-TRP for three days at 30°C. Isolated colonies were selected and spread on DO+5-FOA media (5-Fluoroorotic Acid) to counter-select yeast cells with pTET-HA-AC40sp plasmid. Finally, sc*RPC40* mutant strains were validated by PCR and western blots.

### Protein expression and purification

IN1 harboring a histidine tag for purification followed by a sso7d tag for solubility was produced as described^19^. *Escherichia coli* Rossetta(DE3) cells transformed with the IN1-expressing plasmid were grown in TB with ampicillin and chloramphenicol and grown at 25°C to an OD600 equal to 0.6. Protein expression was induced by addition of 50 μM IPTG and the cells were grown overnight at 25°C. Cells were harvested at 1100g for 15 min at room temperature, washed with PBS 1X and weighed. Purification of IN1 was performed following described procedures^19^.

For Pol III, *Saccharomyces cerevisiae* strain SC1613, encoding a tandem affinity-purification (TAP) tag at the C-terminus of subunit AC40, was provided by Cellzome AG (Heidelberg, Germany). Pol III was purified as described^20^ with some modifications. About 0.8 kg of cells were suspended in buffer A (250 mM Tris-HCl pH 7.4, 20% glycerol, 250 mM (NH_4_)_2_SO_4_, 1 mM EDTA, 10 mM MgCl_2_, 10 µM ZnCl_2_, 10 mM β-mercaptoethanol) and lysed at 4 °C with glass beads in a BeadBeater (BioSpec). After centrifugation, the supernatant was incubated over night at 4 °C with 6 ml of IgG Sepharose (GE Healtcare), then the resin was washed with 20 column volumes of buffer B (50 mM Tris-HCl pH 7.4, 5% glycerol, 250 mM NaCl, 5 mM MgCl_2_, 10 µM ZnCl_2_, 5 mM DTT) and incubated overnight with tobacco etch virus (TEV) protease to elute the protein by removal of part of the TAP-tag. The TEV eluate was further purified using a Mono-Q (GE Healthcare) with a gradient from buffer B to the same buffer containing 1 M NaCl. Pol III eluted at ∼360 mM NaCl. The protein was concentrated to 7-8 mg/ml, frozen with liquid nitrogen and stored at -80 °C until use.

### Preparation of Pol III-IN1 complexes

For the Pol III-IN1 complex, Pol III and IN1 both at 0.5 µM were incubated in 20 mM HEPES-NaOH, pH 7.5, 170 mM NaCl, 2 mM DTT for 30 min at 4 °C. The complex was crosslinked following described procedures with minor modifications^28^. Briefly, the reaction was started by addition of 0.06%(v/v) of glutaraldehyde and, after 5 min incubation on ice, the remains of the cross-linking agent were quenched by adding 50 mM glycine and incubating 5 min on ice.

To prepare the mismatched transcription bubble, non-template DNA (5’-GCAGCCTAGTTGATCTCATAGCCCATTCCTACTCAGGAGAAGGAGCAGAGCG-3’), template DNA (5’-CGCTCTGCTCCTTCTCCTTTCCTCTCGATGGCTATGAGATCAACTAGGCTGC-3’) and RNA (5’-AUCGAGAGGA-3’) from Microsynth were incubated following described procedures^45^. The Pol III EC was assembled by incubating the transcription scaffold at equimolar amounts with the enzyme, at a final concentration of 0.4 μM, for 1 hour at 20 °C in buffer E (10 mM Hepes pH 7.5, 150 mM NaCl, 5 mM MgCl_2_, 5 mM DTT). The sample was then mixed with 0.6 μM IN1 (final concentration) and incubated for 30 min at 20 °C in buffer E to obtain the Pol III EC bound to IN1.

### Cryo-EM grid preparation and data acquisition

The crosslinked Pol III-IN1 complex was concentrated to 1 µM, then 4 µl were applied to glow-discharged copper 300 mesh C-flat 1.2/1.3 holey carbon grids (Protochips) in the chamber of a FEI Vitrobot at 10 °C and 100 % humidity. The grids were blotted for 3.5 sec with blotting force -5 and vitrified by plunging into liquid ethane cooled with liquid nitrogen. Movies were acquired on a FEI Titan Krios (ThermoFisher) electron microscope at 300 keV using a K3 summit (Gatan) direct electron detector operated in ‘super-resolution’ mode with a physical pixel size of 1.06 Å. A total of 3,321 non-tilted movies and 8,900 movies tilted by 20° with 40 frames each were collected with an accumulated total dose of 42.45 e-/Å^2^.

For the Pol III EC bound to IN1, 3 µl of the sample were supplemented with 8 mM CHAPSO (final concentration), applied to glow-discharged copper 300 mesh Quantifoil R1.2/1.3 grids coated with continuous carbon, and incubated in the chamber of a FEI Vitrobot at 24 °C and 100% humidity for 1 min. The grids were blotted for 4 sec with blotting force 2.5 and vitrified by plunging into liquid ethane cooled down to liquid nitrogen temperature. Data were collected on a FEI Titan Krios electron microscope operated at 300 kV, using a K3 summit (Gatan) direct electron detector. Images were acquired at defocus values varying between -1.0 and -2.8 μm at a pixel size of 1.085 Å. A total of 9,327 movies with 40 frames each were collected with an accumulated total dose of 45.12 e-/Å^2^.

### Cryo-EM data processing

For the Pol III-IN1 complex, movies were aligned and dose-corrected using MotionCor2^46^ as implemented in Relion 3.0^47^. Global contrast transfer function (CTF) parameters were estimated using GCTF^48^. Around 1,000 particles were picked manually to generate reference-free 2D classes that were used for template-based auto-picking after low-pass filtering to 20 Å, followed by correction of local defocus of the individual particles using GCTF. The remaining processing was performed in Relion 3.1 (Fig S1). Three-fold binned particles were subjected to reference-free 3D classification and the best class with 885,560 particles was selected and subjected to another round of reference-free 3D classification. The five best classes with 643,858 particles in total were extracted with a 300 pixel box without binning and subsequently refined, followed by two rounds of CTF refinement^49^ and Bayesian polishing. Final post-processing was performed using automatic masking and B-factor sharpening, resulting in a map at 2.5 Å resolution. Five independent runs of 3D classification were performed using masks for the AC40 subunit, the stalk, the protrusion plus clamp head, C11-Nt and C11-Ct. For each case, the best classes were selected and refined. The AC40 mask yielded a map at 2.6 Å resolution with clear density for TD1, while the C11-Ct mask produced maps at 2.9 and 2.8 Å resolution where this domain is present and absent, respectively.

For the Pol III PCC-IN1 complex, movies were aligned and dose-corrected using MotionCor2 as implemented in Relion 3.1, and their CTF parameters were estimated using CTFFIND4^50^. Approximately 550 particles were picked manually and reference-free 2D classes were generated, five of which were used for template-based autopicking after low-pass filtering to 20 Å. Approximately 2,437,000 particles were automatically selected and extracted with a 300 pixel box using Relion 3.1, also employed for subsequent processing (Fig S2). Three rounds of reference-free 2D classification yielded a stack of 469,620 good-quality particles that were used to generate an initial 3D model employing a reference generated from EMD-3180 map filtered to 60 Å. The resulting map was used as a reference for 3D classification to generate four classes. Two of these classes showing Pol III-like shape were joined together (305,253 particles total) and refined to a resolution of 3.8 Å. The resolution of the map was improved to 3.07 Å using particle polishing and CTF refinement.

### Model building and refinement

The available structure of the Poll III EC (PDB code: 5FJ8) was fitted in the 3D maps using UCSF Chimera^51^ and employed as starting point for model building for Pol III subunits and the nucleic acid scaffold. A homology model of the C11 C-terminal Zn-ribbon was generated with Phyre2^52^ and used as reference for model building, while the C11 linker was manually modelled in Coot^53^. Available structures of Pol III complexed to melted DNA (PDB code: 6EU1) and the Pol III-Maf1 complex (PDB code: 6TUT) were used as reference to improve coordinates of the stalk C17/C25 heterodimer, the C82/C34/C31 heterotrimer and the C37/C53 heterodimer. The nucleic acid scaffold DNA model was adapted from a stalled Pol I elongation complex (PDB code: 6H67). Segments corresponding to IN1 and C53-Nt were manually-built *de novo* based only on our maps. The structures were refined using real-space refinement as implemented in Phenix^54^. Refinement statistics are summarized in Table S1. Figures were prepared using PyMOL (Schrödinger Inc.) and UCSF Chimera^51^.

### RNA elongation assay

The nucleic acid scaffold was prepared by mixing equal amounts of template DNA: 5’-CGTAGCGGTATCGTGGTCGAGCGTGTCCTGGTCTAG-3’, non-template DNA: 5′-CGCTCGACCACGATACCGCTACG-3′ and RNA: 5′-Alexa488-CGACCAGGAC-3′ in 20 mM Tris, pH 7.5, 150 mM KCl, heated to 95 °C and slow-cooled to 4 °C. For RNA elongation, 100 nM Pol III was pre-incubated, when required, with 200 nM IN1 in 20 mM Tris pH 7.5, 150 mM KCl, 5 µM ZnCl_2_, 10 mM DTT for 10 min at room temperature. Then 100 nM of the scaffold was added and incubated for another 10 min. To initiate the reaction, 1 mM of NTPs mix (Invitrogen) and 10 mM MgCl_2_ was added and incubated at 37 °C. The reaction was stopped by adding an equal amount of 2x RNA loading dye (8 M urea, 2x TBE, 0.02% bromophenol blue, 10 % (v/v) glycerol) and heating to 95 °C for 5 min. The samples were loaded onto denaturing 20% polyacrylamide gel containing 7 M urea and visualized with a Fujifilm FLA-3000.

### pGALTy1-*his3AI* transposition assays

To estimate the frequency of retrotransposition of pGALTy1-*his3AI*, four independent transformants of each strain were grown to saturation 2 days at 30°C in liquid SC-URA containing 2% raffinose. Each culture was diluted thousand-fold in liquid SC-URA containing 2% galactose and grown 5 days to saturation at 20°C, which is the optimal temperature for Ty1 retrotransposition. Aliquots of cultures were plated on YEPD (100 μl at 10^−5^) and SC-HIS (100 μl at 10^−2^). Plates were incubated for 3 days at 30°C and colonies counted to determine the fraction of [HIS^+^] prototroph. Retrotransposition frequencies were defined as the mean of at least four experiments, and each one performed with four independent transformants.

### PCR assays for detection of Ty1 integration events

To detect endogenous Ty1 insertions, three colonies of each yeast strain were inoculated overnight in YPD at 30°C. Next day, each culture was diluted thousand-fold in YDP to induce Ty1 retrotransposition for 3 days at 20°C and total genomic DNA was extracted according to classical procedures^55^. For the detection of Ty1-*HIS3* insertions, Ty1-*his3AI* retrotransposition was induced as described in the previous section and total genomic DNA was extracted from yeast cultures grown at 20°C for 5 days. Double-strand DNA (dsDNA) concentration was determined using Qubit™ fluorometric quantification (Thermo Fisher Scientific). Ty1-*HIS3* integration upstream of *tG(GCC)C* (*SUF16*) were amplified with PCR primers O-AL27 and O-AB91 and at the *SEO1* subtelomeric gene with PCR primers O-AL27 and O-AL10. Endogenous Ty1 integration upstream of *tG(GCC)C* (*SUF16*) were amplified with PCR primers O-AB46 and O-AB91 and at the *HXT* subtelomeric genes (*HXT13, HXT15, HXT16* and *HXT17*) with PCR primers O-AB46 and O-ABA27.

PCR reaction consisted of 30 ng of dsDNA (or 75 ng for detection of insertions at *HXT* or *SEO1*), 5 μl Buffer 5×, 0.5 μl dNTP 10 mM, 0.625 μl of each primer at 20 μM, 0,25 μl of Phusion DNA Polymerase (Thermo Scientific) in a 25 μl final volume. Amplification was performed with the following cycling conditions in ProFlex™ PCR System (Life Technologies) cycler: 98°C 2 min, 30× [98°C 10 s, 60°C 30 s, 72°C 1 min], 72°C 5 min, and hold 4°C. PCR products were separated on a 1.5% agarose gel.

### Data availability

The structures and associated data of the Ty1 integrase-Pol III complex in the absence of nucleic acids with focused classification on AC40, on C11-Ct with this domain in the funnel, on C11-Ct with this domain disordered, and in the presence of nucleic acids have been deposited in the Protein Data Bank under accession codes 7Z0H, 7Z30, 7Z31, 7Z2Z and the Electron Microscopy Data Bank under accession codes EMD-14421, EMD-14469, EMD-14470, EMD-14468.

## Supporting information

All supplemental data

## Acknowledgements

We thank Tornero Laboratory members Federico M. Ruiz for helpful advice and Srdja Drakulic for technical assistance. We also acknowledge Ernesto Arias-Palomo at CIB-CSIC and Alistair Siebert and Vinod Vogirala at the Diamond Light Source for support in cryo-EM data collection, and Rafael Núñez at the CIB Electron Microsopy Facility for help during cryo-EM grid preparation.

## Funding

JA, JR, PL and CFT were supported by the Agence Nationale de la Recherche (Grant ANR-17-CE11-0025). Work in CFT lab was supported by MCIN/AEI/ERDF (Grants BFU2017-87397-P, RED2018-102467-T and PID2020-116722GB-I00) and by intramural funding from CSIC (Grant 2020AEP152). APP was funded by a predoctoral fellowship from MCIN/AEI (PRE2018-087012). Work in PL lab was supported by intramural funding from Centre National de la Recherche Scientifique (CNRS), the Université de Paris and the Institut National de la Santé et de la Recherche Médicale (INSERM). AAL was supported by a postdoctoral fellowship from Fondation pour la Recherche Médicale (FRM-SPF20170938755).

## Notes

### Competing Interest Statement

The authors have declared no competing interest.

