## Supplementary material for "Structural basis of Ty1 integrase tethering to RNA polymerase III for targeted retrotransposon integration": All supplemental data

**This PDF file includes:**

Tables S1 to S4

Figures S1 to S7

**Table S1. Cryo-EM data collection and refinement statistics**

| | **Pol III-IN1** | | | **Pol III PCC-IN1** |
|---|---|---|---|---|
| Sample support | C-flat 1.2/1.3<br>300 mesh<br>Holey carbon | | | Quantifoil 1.2/1.3<br>300 mesh<br>Continuous carbon |
| Microscope | Titan Krios | | | Titan Krios |
| Detector | K3 | | | K3 |
| Voltage (kv) | 300 | | | 300 |
| Number of Frames | 40 | | | 40 |
| Dose ($e^-/Å^2$) | 42.45 | | | 45 |
| Exposure time (s) | 3 | | | 3 |
| Pixel size (Å/pixel) | 1.06 | | | 1.085 |
| Number of grids | 1 | | | 1 |
| Days of data collection | 3 | | | 2 |
| Collected micrographs | 12221 | | | 9327 |
| Particles after initial 2D class | Not applicable | | | 469620 |
| Particles after initial 3D class | 885560 | | | Not applicable |
| Map name | Map A | Map B | Map C | Map D |
| Final number of particles | 226817 | 143021 | 273119 | 305253 |
| Resolution (Å) | 2.65 | 2.92 | 2.76 | 3.07 |
| AccuracyRotations (°) | 0.597 | 0.733 | 0.714 | 0.631 |
| AccuracyTranslations (pixel) | 0.322 | 0.507 | 0.466 | 0.446 |
| Sharpening B-factor ($Å^2$ ) | -49.3 | -62.1 | -60.2 | -73.5 |
| Ramachandran plot | | | | |
| Outliers (%) | 0.00 | 0.00 | 0.00 | 0.00 |
| Allowed (%) | 7.17 | 7.04 | 5.13 | 7.60 |
| Favoured (%) | 93.83 | 92.96 | 94.27 | 92.40 |
| Map CC (around atoms) | 0.76 | 0.77 | 0.76 | 0.76 |
| RMSD bond lengths (Å) | 0.005 | 0.005 | 0.003 | 0.003 |
| RMSD bond angles (°) | 0.775 | 0.749 | 0.651 | 0.625 |
| All-atom clashscore | 12.49 | 12.83 | 10.87 | 9.09 |
| Rotamer outliers (%) | 0.93 | 0.74 | 0.61 | 0.46 |
| C-beta deviations | 0.04 | 0.02 | 0.00 | 0.06 |
| FSC(model-map) = 0.5 | 2.85 | 2.81 | 2.79 | 3.26 |
| EMDB code | 14421 | 14469 | 14470 | 14468 |
| PDB code | 7Z0H | 7Z30 | 7Z31 | 7Z2Z |

**Table S2. Strains**

| **Strain** | **Genotype** | **Origin** |
|---|---|---|
| LV174 | *MATα ura3Δ851 trp1Δ63 his3Δ200 spt3-101 Δrad52::TRP1* | Ref. 16 |
| PJ69-4A | *MATα trp1-901 leu2-3,112 ura3-52 his3Δ200 gal4Δ gal80Δ LYS2::GAL1-HIS3 GAL2-ADE2 met2::GAL7-lacZ* | Ref. 56 |
| LV1635 | *MATα ura3Δ851 trp1Δ63 his3Δ200 RPC40::HygroB + pTET-HA-RPC40Sc (TRP)* | This study |
| LV1636 | *MATα ura3Δ851 trp1Δ63 his3Δ200 RPC40::HygroB + pTET-HA-RPC40Sp (TRP)* | This study |
| LV1690 | *MATα ura3Δ851 trp1Δ63 his3Δ200 RPC40::HygroB + pTET-HA-RPC40Sp (URA)* | This study |
| LV1857 | *MATα ura3Δ851 trp1Δ63 his3Δ200 RPC40::HygroB + pTET-HA-RPC40Sc* $F_{315}$*A (TRP)* | This study |
| LV1859 | *MATα ura3Δ851 trp1Δ63 his3Δ200 RPC40::HygroB + pTET-HA-RPC40Sc* $F_{315}$*E (TRP)* | This study |
| LV1863 | *MATα ura3Δ851 trp1Δ63 his3Δ200 RPC40::HygroB + pTET-HA-RPC40Sc* $D_{111}$*A (TRP)* | This study |
| LV1873 | *MATα ura3Δ851 trp1Δ63 his3Δ200 RPC40::HygroB + pTET-HA-RPC40Sc* $E_{312}$*A (TRP)* | This study |
| LV1869 | *MATα ura3Δ851 trp1Δ63 his3Δ200 RPC40::HygroB + pTET-HA-RPC40Sc* $I_{320}$*E (TRP)* | This study |
| LV1871 | *MATα ura3Δ851 trp1Δ63 his3Δ200 RPC40::HygroB + pTET-HA-RPC40Sc* $I_{320}$*W (TRP)* | This study |
| LV1897 | *MATα ura3Δ851 trp1Δ63 his3Δ200 RPC40::HygroB + pTET-HA-RPC40Sc* $K_{316}$*E (TRP)* | This study |

**Table S3. Plasmids**

| **Name** | **Vector** | **Genotype** | **Origin** |
|---|---|---|---|
| pAT36 | pAS2ΔΔ | 2µ *AmpR TRP1 GBD-AC40* | Ref. 17 |
| pPL2 | pGAL1Ty1his3AI | 2µ *AmpR URA3 GAL1p*-Ty1*his3*AI | Ref. 30 |
| pCG1 | pGAL1Ty1his3AI | 2µ *AmpR URA3 GAL1p*- Ty1*his3*AI (IN-K$_{617}$A) | Ref. 16 |
| pCG2 | pGAL1-Ty1his3AI | 2µ *AmpR URA3 GAL1p*- Ty1*his3*AI (IN-M$_{619}$A) | Ref. 16 |
| pCG3 | pGAL1-Ty1his3AI | 2µ AmpR *URA3 GAL1p*- Ty1*his3*AI (IN-R$_{620}$A) | Ref. 16 |
| pCG4 | pGAL1-Ty1his3AI | 2µ *AmpR URA3 GAL1p*- Ty1*his3*AI (IN-S$_{621}$A) | Ref. 16 |
| pCG5 | pGAL1-Ty1his3AI | 2µ *AmpR URA3 GAL1p*-Ty1*his3*AI (IN-L$_{622}$A) | Ref. 16 |
| pAL1 | pACTII | 2µ *AmpR LEU2 GAD-IN5*$_{\Delta TD+bNLS}$ | Ref. 16 |
| pAL10 | pACTII | 2µ *AmpR LEU2 GAD-IN5*$_{\Delta TD+bNLS}$ *K*$_{617}$*A* | Ref. 16 |
| pNP67 | pACTII | 2µ *AmpR LEU2 GAD-IN5*$_{\Delta TD+bNLS}$ *W*$_{614}$*A* | This study |
| pRM2 | pCM185 | CEN *AmpR TRP1 pTET-HA-RPC40* | Ref. 17 |
| pRM3 | pCM185 | *CEN AmpR TRP1 pTET-HA-RPC40Sp* | Ref. 17 |
| pAT30 | pCM189 | *CEN AmpR URA3 pTET-HA-RPC40Sp* | Ref. 17 |
| pNP52 | pCM185 | CEN *AmpR TRP1 pTET-HA-RPC40 D*$_{111}$*A* | This study |
| pNP54 | pCM185 | CEN *AmpR TRP1 pTET-HA-RPC40 E*$_{312}$*A* | This study |
| pNP57 | pCM185 | CEN *AmpR TRP1 pTET-HA-RPC40 F*$_{315}$*A* | This study |
| pNP58 | pCM185 | CEN *AmpR TRP1 pTET-HA-RPC40 F*$_{315}$*E* | This study |
| pNP60 | pCM185 | CEN *AmpR TRP1 pTET-HA-RPC40 I*$_{320}$*E* | This study |
| pNP61 | pCM185 | CEN *AmpR TRP1 pTET-HA-RPC40 I*$_{320}$*W* | This study |
| pNP69 | pCM185 | CEN *AmpR TRP1 pTET-HA-RPC40 K*$_{316}$*E* | This study |
| pNP71 | pGAL1-Ty1his3AI | 2µ *AmpR URA3 GAL1p*- Ty1*his3*AI (IN-W$_{614}$A) | This study |

**Table S4. Primers**

| Primer | Sequence 5'- 3' | Gene/Mutation /Plasmid | Reference |
|---|---|---|---|
| O-AL112 | AAATACTAAGAATATGCGTAGTTTAG | pGAD-IN5$_{\Delta TS5+NLS1}$ and pGAL1p-Ty1*his3*AI | This Study |
| O-AL113 | GCTGTGTCTCGTGATACCTTAATTTC | | |
| O-AL114 | AATGCTCACATGGGTTGATAGTAATTTG | pTET-HA-AC40sc D$_{111}$A | |
| O-AL115 | GCAGGGTCAACTTTTAATGGAACCAAG | | |
| O-AL141 | GACACCAGAAGCGATTTTTTCAAATCC GTCAG | pTET-HA-AC40sc E$_{312}$A | |
| O-AL143 | ATGGCACCAGCGCTTTCT | | |
| O-AL123 | TAAATCCGTCAGGATTTTAAAG | pTET-HA-AC40sc F$_{315}$A | |
| O-AL124 | GCAAAAATTTCTTCTGGTGTCATG | | |
| O-AL125 | CAAATCCGTCAGGATTTTAAAG | pTET-HA-AC40sc F$_{315}$I | |
| O-AL134 | ATAAAAATTTCTTCTGGTGTCATG | | |
| O-AL130 | GTTAAAGAATAAGGCTGAGTATTTG | pTET-HA-AC40sc I$_{320}$W | |
| O-AL131 | CACCTGACGGATTTGAAAAAAATTTC | | |
| O-AL132 | ATTAAAGAATAAGGCTGAGTATTTG | pTET-HA-AC40sc I$_{320}$E | |
| O-AL133 | TCCCTGACGGATTTGAAAAAAATTTC | | |
| O-AL169 | GTTGAAGGCAGAATGCATTTTAG | pTET-HA-AC40sc K$_{316}$E | |
| O-AL170 | GAAATTTCTTGACCAGGC | | |
| O-AB46 | GTGATGACAAAACCTCTTCCG | TYB OUT-2 | Ref. 57 |
| O-AB91 | TTTTAGAGTGACACCATCGTAC | SUF16 (SNR33 OUT) | |
| O-ABA27 | GACATGGGCCCCTGTTGCTTATATTGT | HXT13, HXT15, HXT16 and HXT17 | Ref. 18 |
| O-AL10 | CGTTGGTGCTGCATATGTCA | SEO1 | Ref. 16 |
| O-AL27 | GCAAGAGAGATCTCCTACTTTC | HIS3 | |

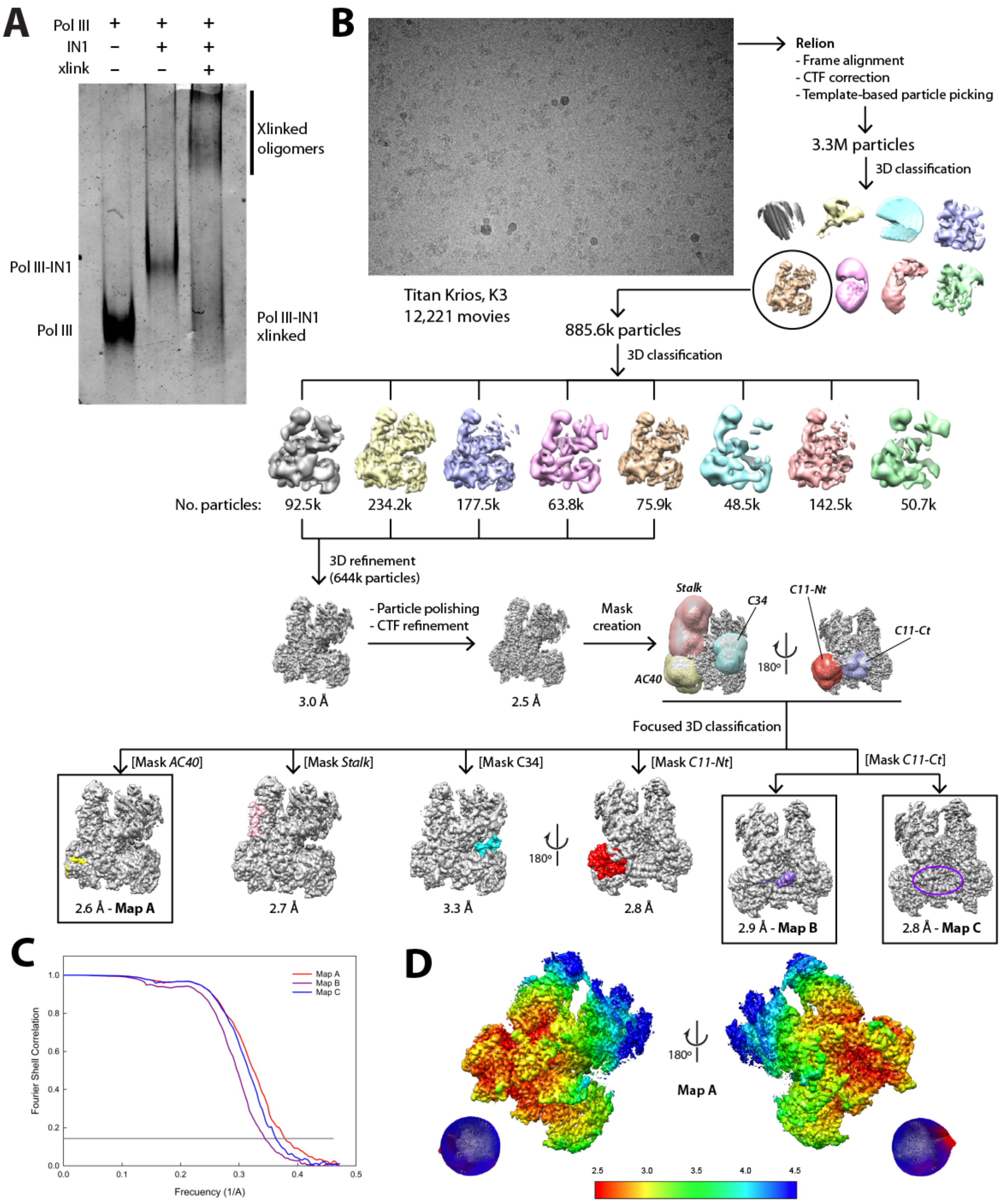


Figure S1. Cryo-EM of the Pol III-IN1 complex. (A) Tris-glycine gel showing differential mobility of free Pol III, the Pol III-IN1 complex, and the latter after crosslinking (xlink). (B) Processing pipeline of the Pol III-IN1 dataset including a representative micrograph. Note that 2D classification was skipped and initial selection of good particles was performed using 3D classification exclusively. (C) FSC curves of maps A, B and C showing a final average resolution of 2.6, 2.9 and 2.8 Å (FSC = 0.143). (D) Local resolution estimation of map A and corresponding angular distribution plot. The two views are related by a 180º rotation around a vertical axis.

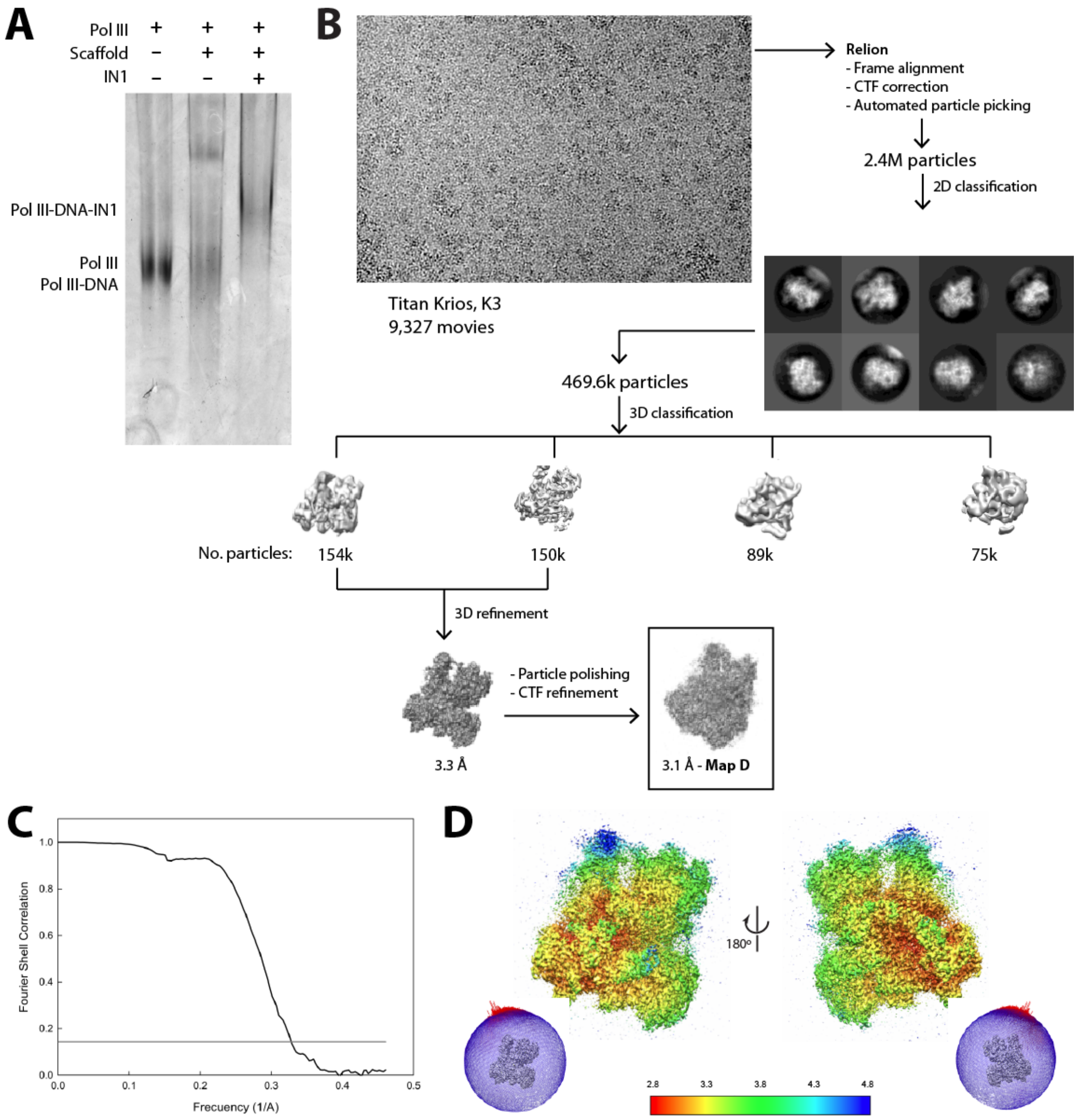


Figure S2. Cryo-EM processing strategy of the complex between Pol III EC and IN1. (A) 4% Tris-glycine gel showing differential mobility of free Pol III, the Pol III EC, and the latter after incubation with IN1. (B) Processing pipeline of the Pol III EC-IN1 dataset including a representative micrograph. (C) FSC curve of the map derived from the best group of particles, showing a final average resolution of 3.1 Å (FSC = 0.143). (D) Local resolution estimation of the map and corresponding angular distribution plot. The two views are related by a 180º rotation around a vertical axis.

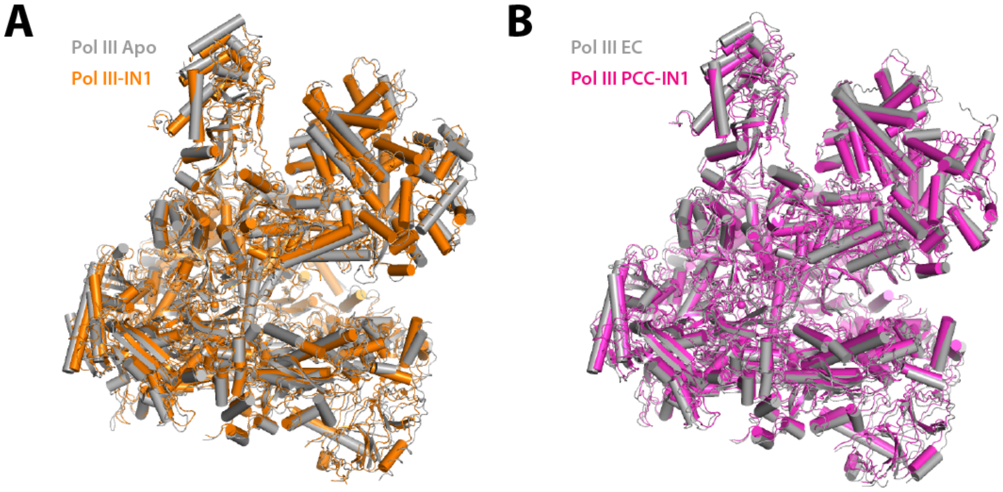


Figure S3. Structural comparison of Pol III-IN1 complexes with reported structures. (A) Superposition of the Pol III-IN1 structure (orange) and that of free Pol III in the closed conformation (PDB 5FJ9, gray), with root-mean-square-deviation (RMSD) of 2.9 Å over 35,981 atoms. (B) Superposition of the Pol III PCC-IN1 structure (magenta) and that of the canonical Pol III-EC (PDB 5FJ8, gray) with RMSD of 1.9 Å over 34,4414 atoms.

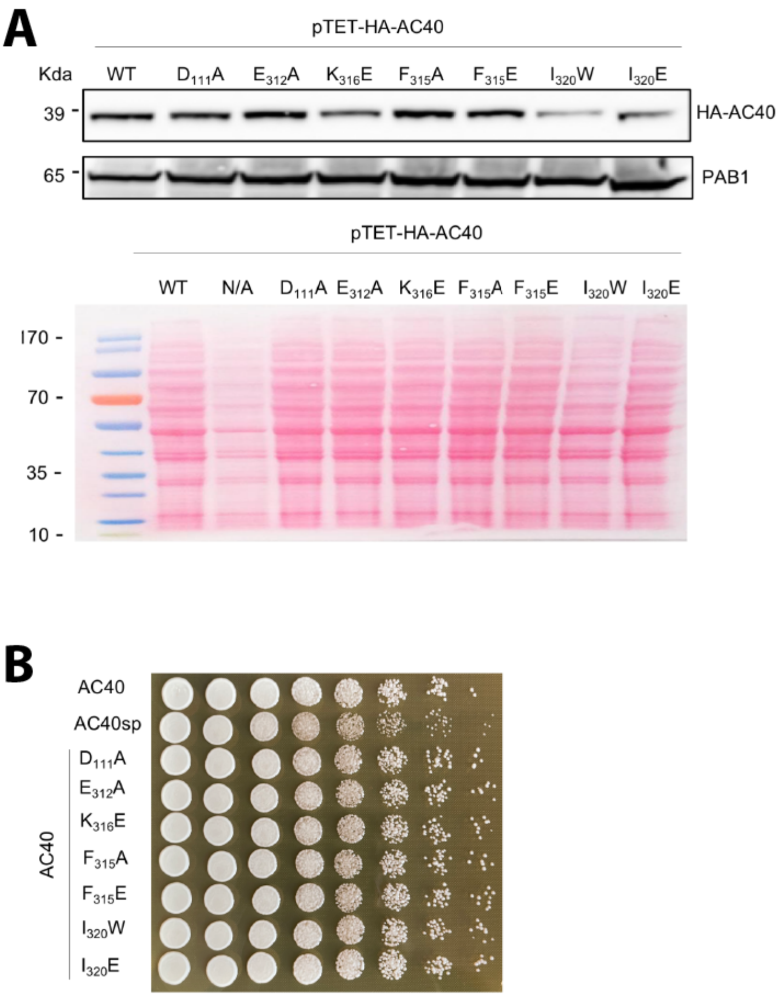


**C**

| Strain genotype | Ty1-his3AI retromobility ± SD (x10-3) | Fold activation | P-value (Welsch t-test) |
|---|---|---|---|
| AC40 | 6.03 ± 0.85 | 1 | N/A |
| AC40sp | 3.26 ± 0.53 | 0.54 | 0.0134 (*) |
| AC40 $D_{111}A$ | 3.40 ± 0.89 | 0.56 | 0.0208 (**) |
| AC40 $E_{312}A$ | 3.09 ± 0.13 | 0.51 | 0.0246 (**) |
| AC40 $K_{316}E$ | 3.74 ± 0.73 | 0.62 | 0.0250 (**) |
| AC40 $F_{315}A$ | 6.58 ± 1.20 | 1.09 | 0.5589 (ns) |
| AC40 $F_{315}E$ | 3.55 ± 0.83 | 0.59 | 0.0227 (**) |
| AC40 $I_{320}W$ | 3.64 ± 0.76 | 0.60 | 0.0226 (**) |
| AC40 $I_{320}E$ | 4.06 ± 0.92 | 0.67 | 0.0528 (*) |

Figure S4. Control experiments related to AC40 mutant analyses. (A) Expression of HA-AC40 mutant proteins. Whole cell extracts of the indicated strains analyzed by western blotting using anti-HA antibodies, revealing HA-AC40 WT and mutant proteins (top panel). Total proteins were detected with the reversible Ponceau staining method (bottom panel) and molecular weights are indicated (kDa). Whole-cell extract samples were prepared for immunoblotting from 10 $OD_{600}$ of the cell culture by TCA precipitation. (B) Growth comparison of WT and mutant *RPC40* strains. 1 $OD_{600}$ of overnight cell culture was used to make 5-fold serial dilutions, starting from $10^0$. Cells were dropped on YPD plates and incubated 48h at 30°C. (C) Retrotransposition frequency of p*GAL1*-Ty1*his3AI* in WT and mutant *RPC40* strains. Values are mean ± SD, $n = 3$ experiments, each performed with four independent colonies. Two-sided Welch's t test was used allowing unequal variance. * $p < 0.05$; ** $p < 0.01$; *** $p < 0.001$; **** $p < 0.001$; ns, not significant. Statistical tests were performed using GraphPad Prism version 9.0.0. N/A is a non-relevant sample in this experiment, whose corresponding lane has been cut from the Western blot for the final set-up.

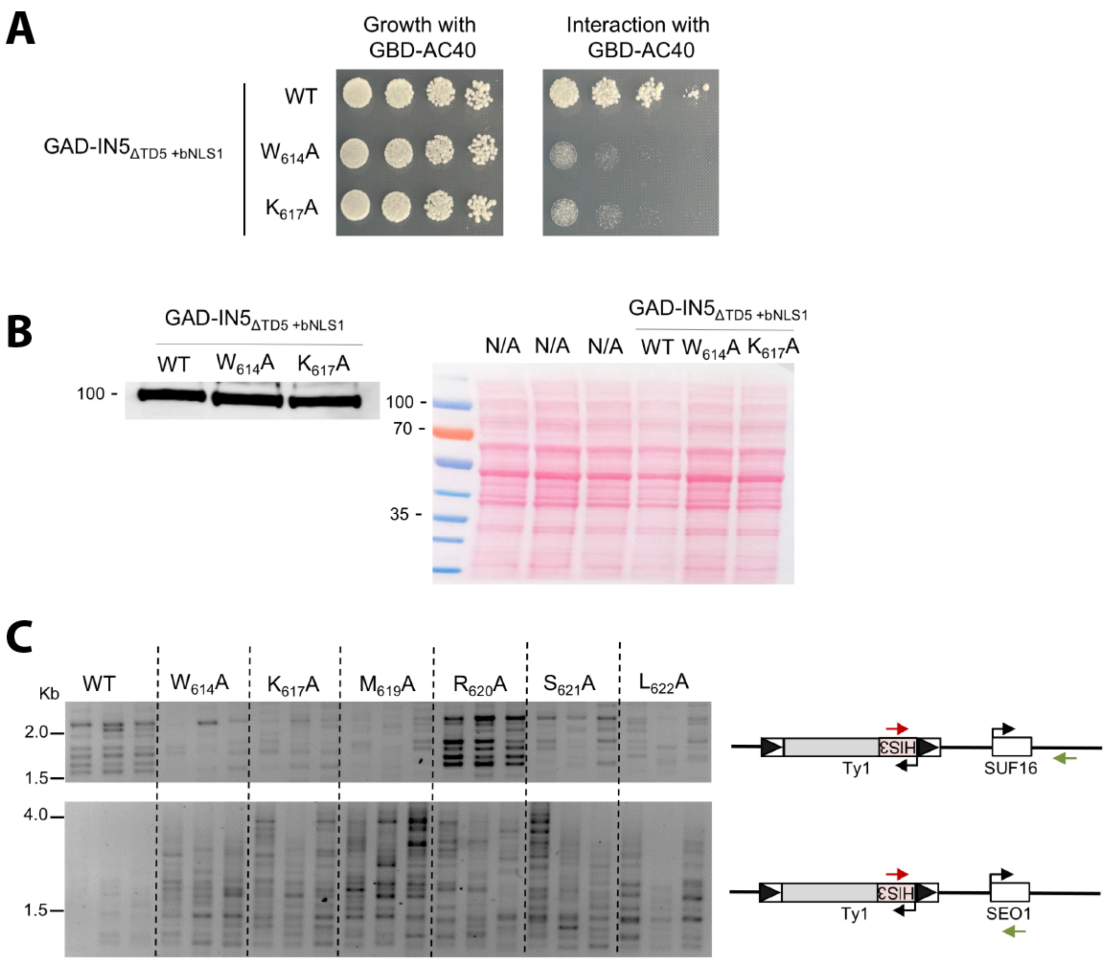


**D**

| pGAL-Ty1his3AI | Ty1-his3AI retromobility ± SD (x10-3) | Fold activation | P-value (Welsch t-test) |
|---|---|---|---|
| **WT** | 5.25 ± 0.81 | 1 | N/A |
| **W614A** | 4.23 ± 0.70 | 0.80 | 0.176432 (ns) |

---

<u>Figure S5. Characterization of Ty1 integrase W614A mutant.</u> (A) Two-hybrid interaction between GBD-AC40 and WT or mutant GAD-IN5$_{\Delta TD5+bNLS}$ in strain PJ69-4A, which contains the *HIS3* reporter to detect positive interactions (James et al., 1996). GAD-IN5$_{\Delta TD5+bNLS}$ harbors the integrase sequence of Ty1 that interacts with AC40 in the integrase sequence of Ty5, in place of the targeting domain of Ty5, and is good indicator of interaction with AC40 (Asif-Laidin et al., 2020). Cultures of transformants were grown overnight at 30°C in synthetic complete medium lacking leucine and tryptophan (SC-LEU-TRP) to maintain plasmid selection. 5-fold serial dilutions of aliquots of 1 $OD_{600}$, washed in 1 ml of $H_20$, were plated on SC-LEU-TRP (growth control) or SC-LEU-TRP-HIS (interaction), starting from $10^0$. Plates were incubated 2 days at 30°C. The assay is a representative example of at least two biological replicates. The $K_{617}A$ mutant was used as control for loss of interaction between IN1 and AC40 (Asif-Laidin et al., 2020). (B) Expression of WT and mutant GAD-IN5$_{\Delta TD5+bNLS}$ proteins. Whole cell extracts of the indicated strains analyzed by western blotting using anti-GAD antibodies (*left panel*). Total proteins were detected with the reversible Ponceau staining method (*right panel*) and molecular weights are indicated (kDa). Whole-cell extract samples were prepared from 1 $OD_{600}$ of the cell culture by TCA precipitation. N/A are non-relevant sample in

this experiment. (C) Detection of Ty1-*HIS3 de novo* insertions generated in cells transformed by a plasmid expressing WT or mutant Ty1*his3AI* elements, bearing the indicated substitutions of conserved residues, from the *GAL1* promoter. Insertions upstream of the *SUF16* and *SEO1* genes (Pol III-transcribed and subtelomeric genes, respectively) were detected by PCR using a primer in *HIS3* (red arrow) and a primer in the locus of interest (green arrow). Ty1 retrotransposition was induced by growing cells 3 days at 20°C. Total genomic DNA was extracted from three independent cultures. (D) Retrotransposition frequency of WT and $W_{614}A$ p*GAL1*-Ty1*his3AI* mutant in a *spt3-101 rad52*Δ strain (LV174) to avoid both trans-complementation of the mutant IN1 by endogenous WT IN1 and Rad52-dependent recombination events (Asif-Laidin et al., 2020). Values are mean ± SD, $n = 3$ experiments, each performed with four independent colonies. Two-sided Welch's t test was used allowing unequal variance. ns, not significant. Statistical test was performed using GraphPad Prism version 9.0.0.

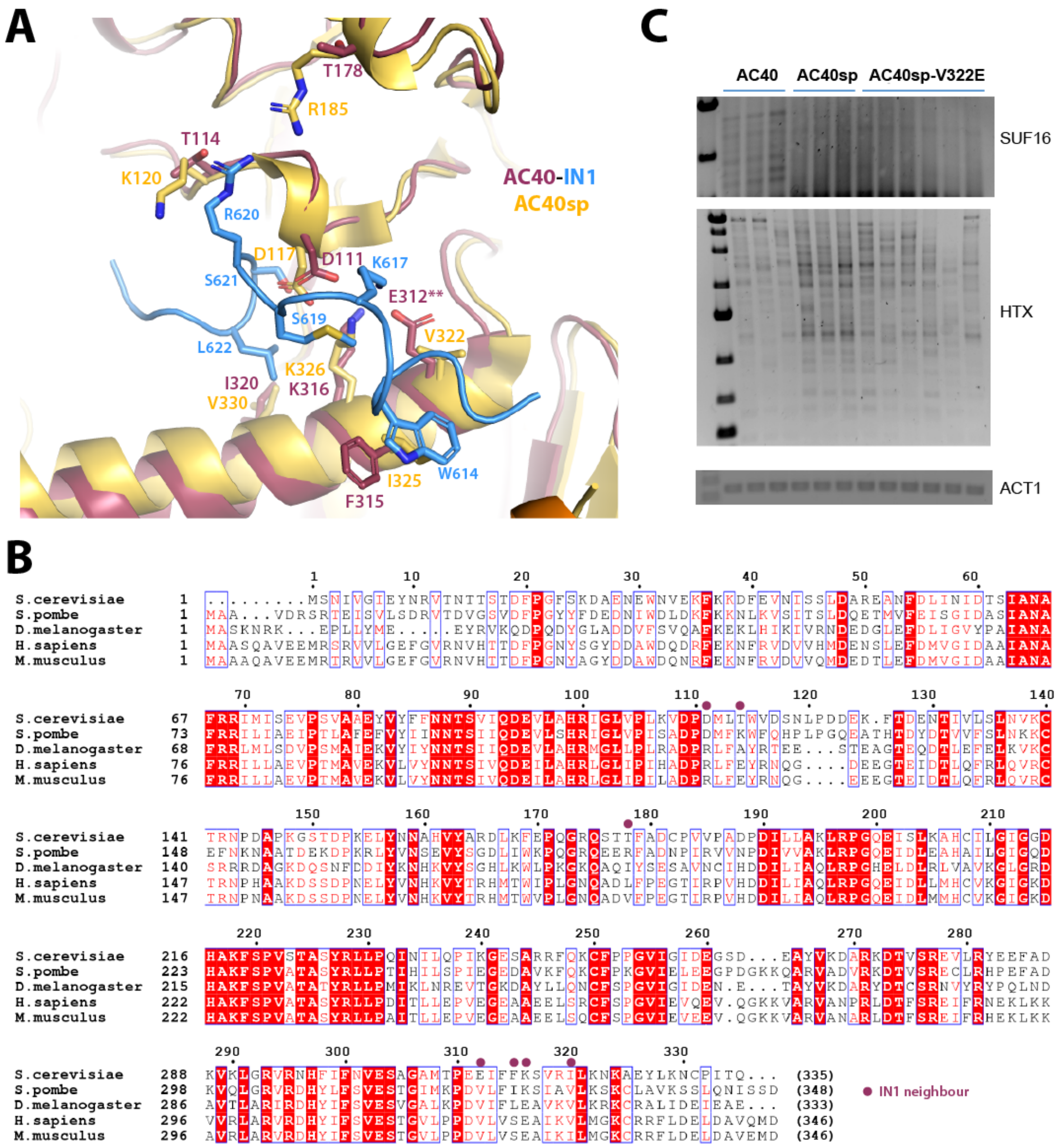


<u>Figure S6. Comparison with *S. pombe* AC40.</u> (A) Structural comparison between *S. cerevisiae* AC40 from Pol III-IN1 (dark red) and *S. pombe* AC40 (AC40sp) from available *S. pombe* RNA polymerase I coordinates (PDB 7AOE, yellow). TD1 is shown in blue. (B) Alignment of AC40 subunits from different organisms, where a dark purple circle indicates residues in direct contact with IN1 that have been mutated in this study. (C) Detection of endogenous Ty1 insertions upstream of the *SUF16* and *HXT* subtelomeric genes (*HXT13*, *HXT15*, *HXT16* and *HXT17*) by PCR in strains expressing AC40, AC40sp or AC40sp harboring glutamate substitution of V322. Ty1 retrotransposition was induced during 3 days in YPD media. Total genomic DNA was extracted from three independent cultures. *ACT1* is genomic DNA quality control.

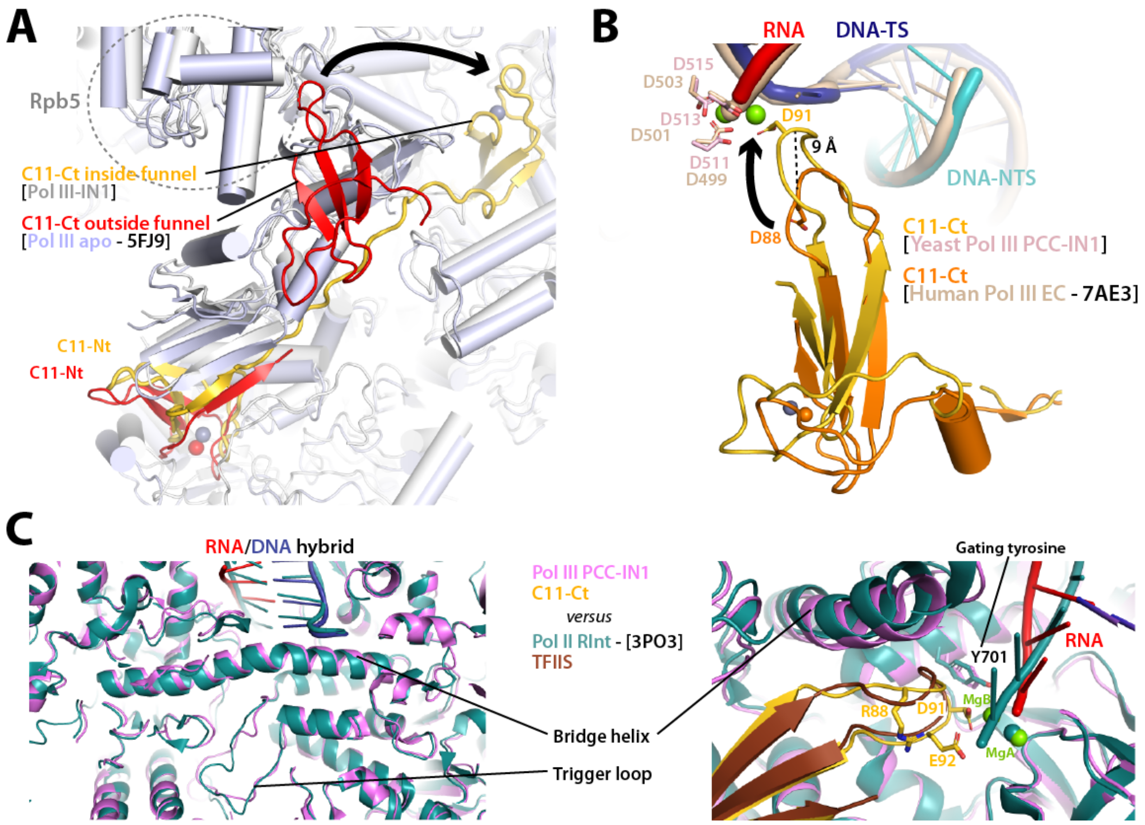


Figure S7. Structural comparisons around subunit C11. (A) Structural comparison of Pol III-IN1 (grey) with free Pol III (PDB 5FJ9, light blue). C11 in these structures is in yellow and red, respectively. (B) Comparison of the yeast Pol III EC-IN1 structure, representing a post-cleavage complex (PCC) of RNA incision (pink), with human Pol III EC (PDB 7AE3, light brown). Nucleic acids in Pol III PCC-IN1 are cyan, blue and red for the non-template strand (NTS) of DNA, template strand (TS) of DNA and RNA, respectively. (C) Structural comparison of Pol III PCC-IN1 (magenta) with Pol II-TFIIS in the rescue intermediate complex (RInt; PDB 3PO3, teal), with close-up views around the bridge helix (left panel) and C11-Ct (right panel).
